# Ion-water coupling controls class A GPCR signal transduction pathways

**DOI:** 10.1101/2020.08.28.271510

**Authors:** Neil J. Thomson, Owen N. Vickery, Callum M. Ives, Ulrich Zachariae

## Abstract

G-protein-coupled receptors (GPCRs) transmit signals across the cell membrane, forming the largest family of membrane proteins in humans. Most GPCRs activate through an evolutionarily conserved mechanism, which involves reorientation of helices and key residues, rearrangement of a hydrogen bonding network mediated by water molecules, and the expulsion of a sodium ion from a protonatable binding site. However, how these components interplay to engage the signal effector binding site remains elusive. Here, we applied information theory to molecular dynamics simulations of pharmaceutically important GPCRs to trace concerted conformational variations across the receptors. We discovered a conserved communication pathway that includes protein residues and cofactors and enables the exchange of information between the extracellular sodium binding site and the intracellular G-protein binding region, coupling the most highly conserved protonatable residues at long distance. Reorientation of internal water molecules was found to be essential for signal transmission along this pathway. By inhibiting protonation, sodium decoupled this connectivity, identifying the ion as a master switch that determines the receptors’ ability to move towards active conformations.

## Introduction

G-protein coupled receptors (GPCRs) form the largest family of cell surface receptor proteins in humans with over 800 members. Spanning the plasma membrane, GPCRs act as signal transducers to enable transmembrane communication into the cell. Extracellular ligand binding leads to conformational changes in the receptors that expose effector protein binding sites on their intracellular face. The effectors, including hetero-trimeric G-proteins and *β*-arrestins, initiate a spectrum of intracellular signalling cascades that cause a variety of physiological changes (1). As such, GPCRs form the main target for drug therapies, with over 30% of US food and drug administration (FDA) approved drugs targeting about 108 different GPCRs (2).

Class A receptors comprise the vast majority of GPCRs. High-resolution crystal structures of class A GPCRs in the inactive state show a sodium (Na^+^) ion bound to Asp^2.50^ (Ballesteros-Weinstein numbering system), a highly conserved, ionizable residue central to their transmembrane domain (Fig. 1). By contrast, structures of receptor active states reveal a collapsed sodium binding site, unable to accommodate the ion (3, 4). Consequently, a wide range of studies have suggested a role for sodium in GPCR activation (4– 10). The absence of sodium is likely to trigger protonation of Asp^2.50^ (6, 11, 12), while the sodium binding site is further embedded in a conserved network of water molecules that connect polar residues and undergo significant re-positioning upon activation (13–15).

**Fig. 1.**
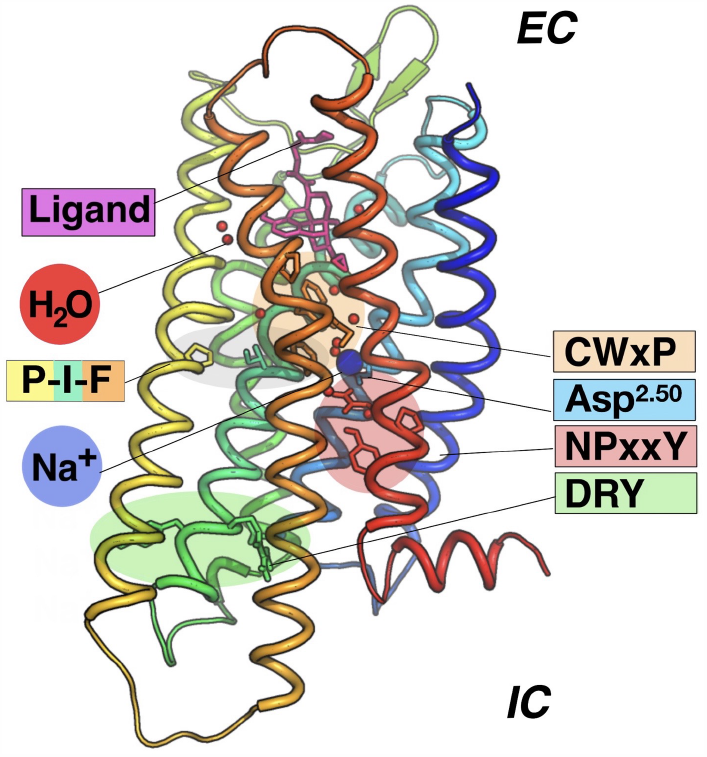
Key class A GPCR features and microswitches. Shown are the ligand binding pocket, internal waters, sodium ion (Na^+^), conserved microswitch motifs, and the primary sodium binding residue Asp^2.50^ in the *µ*-opioid receptor (pdb: 4dkl) as example. EC and IC denote the extracellular and intracellular side, respectively. Important microswitch motifs include the CWxP motif at the ligand binding site, linked directly to the sodium ion binding pocket, and the P-I-F motif, which is strongly coupled to the presence of ligands (23). The NPxxY^7.53^ motif is located proximal to the ion binding site and includes Tyr^7.53^, which can gate a transmembrane water channel according to previous simulation studies (12, 15).

Transitions between distinct rotameric conformations of evolutionarily conserved residues, termed microswitches, are believed to underpin the large scale conformational changes of transmembrane helices that govern activation (1, 13, 16–23). Many microswitches form part of the interconnected polar network. Amongst these residues, the ionizable DRY motif at the intracellular face plays a key role in G-protein binding, and forms ionic locks that maintain the inactive receptor state (1, 20, 24) (Fig. 1).

These observations suggest that the ion binding site, protonation, water-mediated interactions, and conserved residues act together to affect G-protein signalling. However so far, the functional interaction of these elements remains insufficiently understood (4, 14, 25). Here, we hypothesized that these components communicate state changes with each other to induce the global conformational changes determining receptor activation.

GPCR signal transduction is a process whereby information (i.e. about a ligand binding event) is sent, encoded, transmitted and received, in parallel with the tenets of an information system (26). Therefore, Na^+^, water, and microswitches can be seen to act as essential elements of an information system. The function of water molecules in GPCRs has so far predominantly been studied from a structural perspective (14, 27). To resolve their role in signalling, we applied Shannon’s mutual information (26, 28) and McGill’s interaction information (also called co-information) (29–32), as “State Specific Information” (SSI), in order to quantify information shared through coupled transitions of discrete residue and internal water states. In contrast to information theoretic approaches that analyze correlations in protein dynamics in cartesian space (most often C_*α*_ motions) (6, 31, 33– 37), SSI derives from internal coordinate and state-based mutual information frameworks (38–41). It is therefore closely aligned with the concept of GPCR microswitches by requiring a change between at least two distinct conformational states for information exchange.

By combining two independent molecular dynamics simulations with ensembles which differ only in the condition of a specific component, e.g. whether Asp^2.50^ binds a sodium ion, the combined ensemble reflect the effects of that change (Fig. 2). Similar to the Jensen-Shannon divergence (42, 43), SSI then quantifies information shared from that components change in state caused by the transition between individual ensembles and the resulting coupled changes in internal residues and water states in the combined ensemble. As a consequence, SSI establishes causality by tracing the information transfer back to a root cause (see also Materials and Methods).

**Fig. 2.**
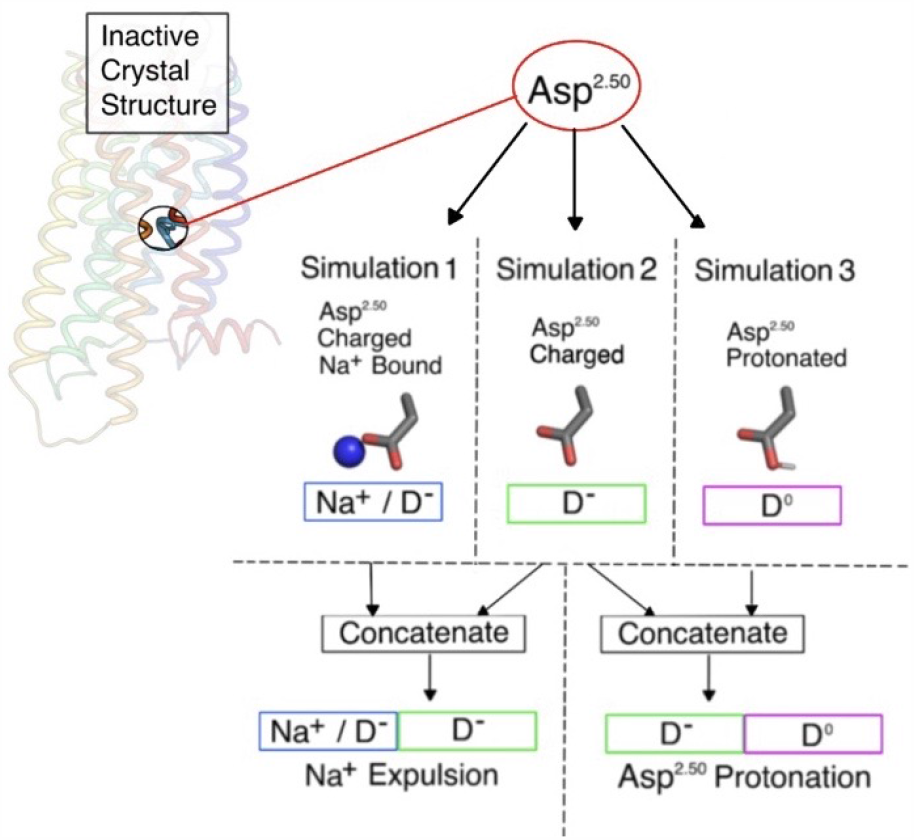
Molecular dynamics simulation approach. For each of the three class A GPCRs, three independent simulations with unique Asp^2.50^ and ion binding states were initiated from the inactive crystal structure: Asp^2.50^ charged with sodium bound (Na^+^/D^-^), Asp^2.50^ charged without sodium bound (D^-^), and Asp^2.50^ protonated without sodium (D^0^). Receptor transitions caused by expelling sodium, and subsequently protonating Asp^2.50^, were represented by concatenating two independent trajectories into one large trajectory. This reflects a binary change to the state of the focal site, corresponding to 1 bit of information (e.g. ‘protonated or de-protonated’), which acts as the source of information.

Accordingly, we conducted all-atom molecular dynamics simulations initiated from three inactive-state, antagonist-bound class A GPCRs: the A_2A_-adenosine receptor (A_2A_AR), the *µ*-opioid receptor (*µ*OR), and the *δ*-opioid receptor (*δ*OR). For each receptor, independent molecular dynamics simulation data of 1.7 *µ*s length was collected, performed in three different receptor states of the main ion binding residue: Asp^2.50^ in a charged state with sodium bound (Na^+^/D^-^), Asp^2.50^ in a charged state with sodium removed (D^-^), and Asp^2.50^ in a protonated state without sodium (D^0^). Transitions between states were represented by pairwise combinations of independent simulations in the respective states (Fig. 2) and analyzed using SSI. Thereby, we extracted information from persistent changes in internal residue and water states occurring in all three receptor types that are coupled specifically to sodium binding or protonation of Asp^2.50^.

Our results reveal how ion binding and Asp^2.50^-protonation are coupled to the dynamics of internal water molecules and protein microswitch conformations. Altogether, these elements form a long-range information transfer pathway between two highly conserved ionizable regions Asp^2.50^ and the intracellular DRY motif at the G-protein binding site. By determining Asp^2.50^-protonation, sodium binding is found to act as the master switch which inhibits concerted transitions towards active state conformations throughout the receptor.

## Results

We first monitored the water occupancy of protein cavities. Five pockets, located in receptor regions of highly conserved motifs, exhibited large probability densities for internal water in all simulations (Fig. 3A,C). Four of these sites are conserved in class A GPCRs (14). Information transferred across the water sites was calculated based on pocket occupancy as well as water orientation. We then investigated information exchanged by coupled conformational state transitions as SSI between 7875 residue and internal water pairs for each of the three receptors. The sodium binding and protonation site was treated as an independent binary residue microswitch (the information source, see Fig. 2) To elucidate pathways conserved across all three receptors, we determined the geometric mean of results from all common residue Ballesteros-Weinstein positions, totaling 121 residues.

**Fig. 3.**
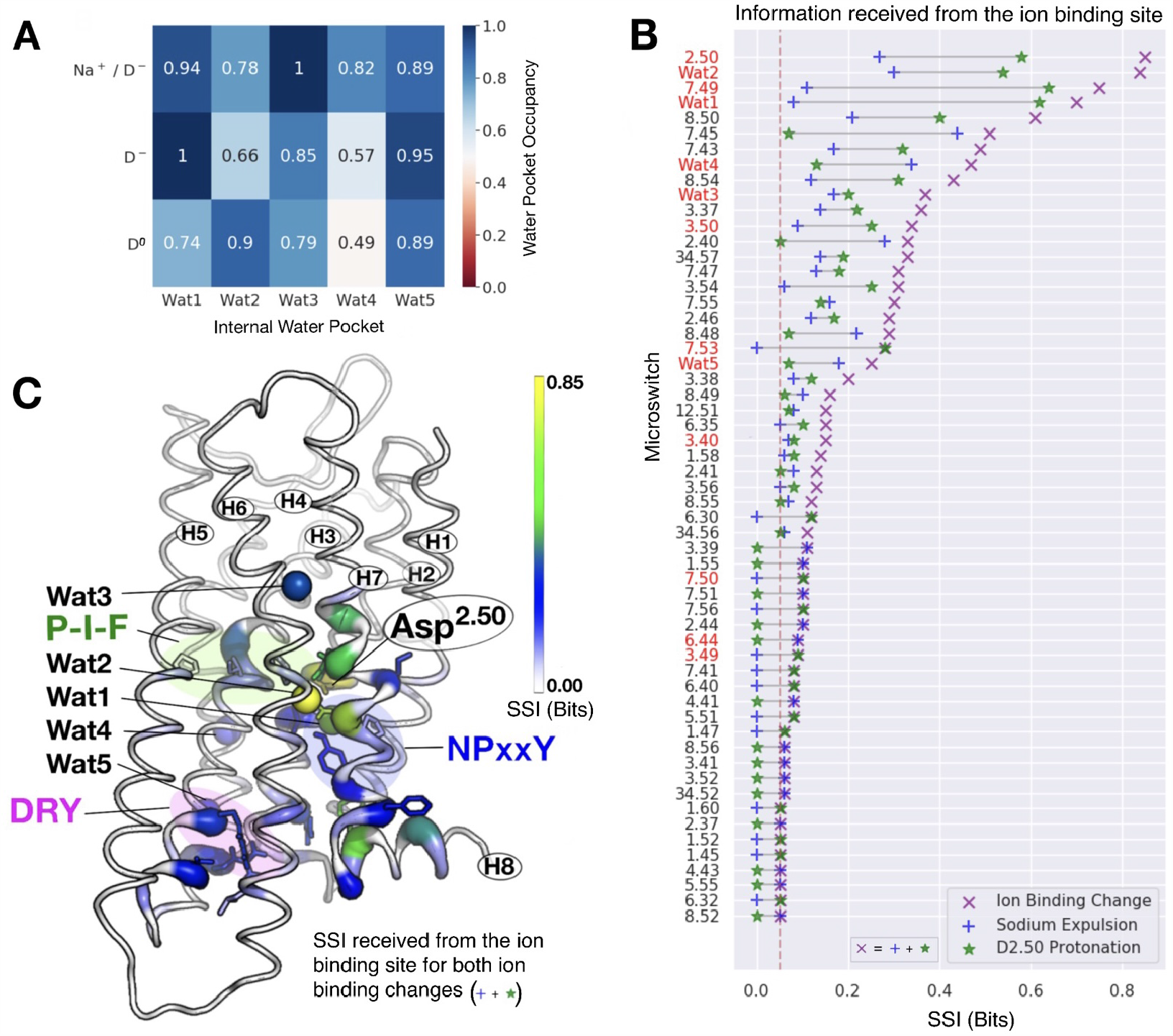
State Specific Information (SSI) reveals information flow propagating through the receptor. **(A)** Water pocket occupancy averaged across all three receptors throughout the simulations; a value of 1 reflects complete occupancy in all receptors over the entire simulated time. **(B)** Information sent from the ion binding site to protein microswitches and internal water molecules (SSI) in the form of coupled conformational transitions. Shown are all significant shared SSI values in bits for sodium expulsion and Asp^2.50^-protonation (each corresponding to 1 bit of source information). The significance threshold is depicted by a vertical red dashed line (0.05 bits), and internal waters, key residues and motifs are highlighted in red. High SSI values reflect a strong coupling between the binary state change of the ion binding site and the conformational state change of a microswitch or water. **(C)** Information flow from the ion binding site projected as colour-code (from white, 0 bits to yellow, 0.85 bits SSI) onto the *µ*OR crystal structure (pdb:4dkl). Key microswitches are shown as sticks, water molecules as spheres. Microswitches involved in SSI transfer form a pathway bridging the ion binding site and the DRY motif at the G-protein binding site.

### An information transfer pathway from the Na^+^ pocket to the intracellular G-protein binding interface

Rather than affecting the entire protein, SSI was shared between a small group of key microswitch and water pairs, both for sodium expulsion and Asp^2.50^-protonation (∼5% and 6%, respectively). Within this core group, our analysis revealed long-range information transfer sent from the ion binding site to the P-I-F motif, the NPxxY motif, and the distal ionizable DRY motif involved in the G-protein binding interface (Fig. 3B). We found that the conformation of residue Arg^3.50^ of the DR^3.50^Y motif, at a distance of more than 20 Å, was strongly impacted by Asp^2.50^-protonation (Fig. 3B).

Internal water molecules acted as important players in the transmission of information emerging from the ion binding site. The conformations of the water molecules were more indicative of the state of the ion binding site than the majority of the protein residues (Fig. 3B). This suggests that water molecules are an integral part of the signal transduction mechanism, and that the internal water molecules form additional elements in an extended set of microswitches. Furthermore, it is known that water 1 (Wat1) forms interactions present in the inactive receptor state, while interactions with water 2 (Wat2) are correlated with the active receptor state (14). In agreement with this, protonation of Asp^2.50^, which has been linked to activation (12), diminished the occupancy of site Wat1 by ∼25%, whereas it increased Wat2 occupancy by ∼25%.

The magnitude of information received from the ion binding site across all three receptors, projected onto the *µ*OR crystal structure, reveals a pathway that includes water and transmits information between Asp^2.50^ and the DRY motif (Fig. 3C). Information travels in the form of propagated conformational changes along helix 7 and across the receptor base. The tight connectivity of coupled transitions on this route allows changes at the ion binding site to affect the conformation of the distal G-protein binding site, establishing long-range coupling between the ionizable residues Asp^2.50^, Asp^3.49^, and Arg^3.50^. The ability of these residues to change protonation state has previously been suggested to be a hallmark of class A GPCR activation (24). Following protonation of Asp^2.50^, the conformational change at Arg^3.50^ also impacted the conformation of residue X^6.30^ (the helix 6 ionic lock), and position X^34.57^ on intracellular loop 2 (ICL2), which is known to play a role in effector protein binding specificity and bias (44–46). Taken together, these results therefore show that the sodium binding pocket communicates at long-range with key interfacial residues that are crucial for effector protein interaction, receptor activation and signal bias.

### Efficient information transfer requires coupling of Asp^2.50^-protonation to water

We found that two key motifs implied in receptor activation, the NPxxY and the DRY motif, became conformationally coupled after Asp^2.50^-protonation (Figs. 3B; for full SSI data see Supplementary Information). Unlike sodium expulsion, Asp^2.50^-protonation caused all residues of the N^7.49^PxxY motif to undergo various degrees of conformational rearrangement. Structurally, Asp^2.50^-protonation directly altered the Asp^2.50^-Asn^7.49^ hydrogen bonding pattern, the conformation of the hydrophobic barrier created by Tyr^7.53^ (12), and the orientation of water molecules Wat1 and Wat2, proximal to the NPxxY motif (Fig. 3B).

To quantify the functional effect of the intercalated water molecules Wat1-Wat5 (Fig. 3A,C) on coupled conformational changes, we calculated the co-information parameter (co-SSI, see Materials and Methods for details) (29–32). Here, positive or negative co-SSI values indicate that the coupling between the ion binding site and each microswitch is stabilised or destabilised, respectively, by their interaction with water molecules. This is equivalent to water amplifying or attenuating the shared information.

We observed that the water network had a particularly pronounced effect on the information pathway upon Asp^2.50^protonation (Fig. 4A,B). Specifically, reorientation of Wat1 and Wat2 was critical to establish communication between the NPxxY and DRY motifs. It also stabilized conformational changes along the pathway to the G-protein binding site as a whole, amplifying the majority of SSI transfer from the protonation site to the microswitches.

**Fig. 4.**
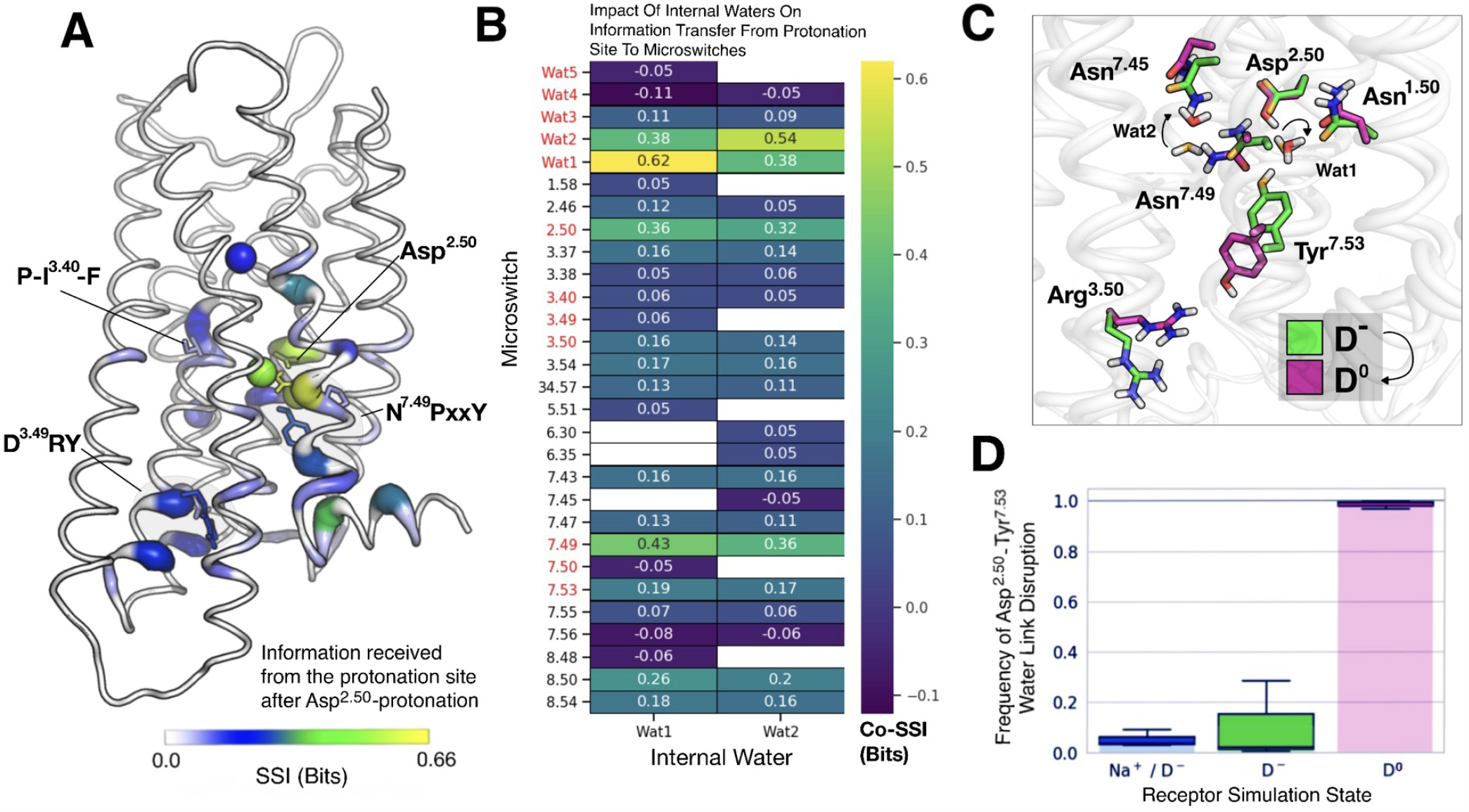
Water molecules are crucial to establish communication between NPxxY and DRY motifs upon Asp^2.50^-protonation. **(A)** Information received by all three receptors after Asp^2.50^-protonation (SSI) projected onto the *µ*OR structure (pdb: 4dkl). **(B)** Impact of the two water molecules, Wat1 and Wat2, on information transfer to the microswitches (co-SSI). Microswitch motifs and internal waters are highlighted in red. Co-SSI equals SSI when two out of three components are identical (e.g., Wat2, Wat2). **(C)** Exemplar frames from molecular dynamics simulations of *µ*OR in states D^-^ (Asp^2.50^ charged, side chain carbon atoms in green and oxygen atoms in orange) and D^0^ (neutral Asp^2.50^, carbon in magenta and oxygen in red). **(D)** Frequency of disruption of the Asp^2.50^–Tyr^7.53^ water bridge, averaged across the three receptors and normalised (box: dataset quartiles; whiskers: maximum and minimum disruption frequency of the three receptors). Asp^2.50^-protonation disrupts the water bridge, enabling coupling between Tyr^7.53^ and Arg^3.50^.

The negative value of co-information between Pro^7.50^, Wat1, and the Asp^2.50^-protonation state, by contrast, implies that reorientation of Wat1 destabilises the conformational state of Pro^7.50^. This may contribute to the kinking of helix 7 seen upon activation (for full co-SSI data see Supplementary Information).

### Molecular mechanism of information transfer propagating from the ion binding site

To elucidate how information transfer is initiated by the protein and water on the molecular level, we next analyzed the major conformational changes caused by Asp^2.50^-protonation. In all simulations of receptors in the Asp^2.50^-deprotonated state (Na^+^/D^-^ and D^-^), Wat1 acted as a H-bond donor with the side chain oxygen atoms of Asp^2.50^ and Asn^1.50^, and as a H-bond acceptor with Tyr^7.53^, where the Tyr OH group pointed towards the Wat1 pocket (Fig. 4C).

By contrast, Asp^2.50^-protonation triggered substantial Hbonding rearrangements in all three receptor types, as Wat1 rotated to accept a H-bond from the protonated Asp^2.50^ side chain. Furthermore, the water bridge between Asp^2.50^ and Tyr^7.53^ was fully disrupted (Fig. 4D). Thereby, Tyr^7.53^ flipped into its downward state in the protonated A_2A_AR and *µ*OR (Fig. 4C). This downward movement has previously been shown to correlate with the opening of a hydrophobic gate upon receptor activation, forming a continuous pore through the receptor (12, 15)). In the *δ*OR, an identical transition was observed upon further protonation of Asp^3.49^ (D^3.49^RY motif), a residue that is coupled to Asp^2.50^-protonation through the information pathway.

Co-SSI indicated that protonation-dependent reorientation of Wat1 was essential for altering the hydrogen bonding pattern that enabled the Tyr^7.53^ downward swing (Fig. 4B), initiating communication with the DRY motif. In the *µ*OR, this resulted in an upward movement of Arg^3.50^ into its active, Gprotein binding conformation (Fig. 4C). In addition, the interhydroxyl distance between Tyr^7.53^ and Tyr^5.58^ was reduced in the *µ*OR and A_2A_AR. In the *µ*OR, Asp^2.50^-protonation was thereby sufficient to decrease the Tyr^7.53^-Tyr^5.58^ interhydroxyl distance to the distribution seen in the active state, when compared to data from a 1.7-*µ*s simulation of the active crystal structure (pdb: 6ddf) (Fig. S3). We also found that the G-protein binding cavity opened upon further protonation of Asp^3.49^ in the *δ*OR, as measured by the distance between Thr^2.39^-C*α* and Ile^6.33^-C*α* (Fig. S4). These results indicate that the protonation of two conserved ionizable residues connected by the information pathway, Asp^2.50^ and Asp^3.49^, may play a direct role in receptor activation and opening of the G-protein binding cleft.

### Sodium restrains key microswitches to a single conformation

The sodium-bound form of class A GPCRs is known to be associated with their inactive state (3, 4, 7, 12). Notably, three residues around the sodium binding site known to form a hydrogen bonding triplet with Wat1, Asp^2.50^, Tyr^7.53^ and Asn^1.50^ (a 100% conserved residue in class A GPCRs), were restrained to highly ordered single conformations in all three sodium-bound receptors. This was reflected by their zero conformational state entropies (see Materials and Methods) (Fig. 5). The water-mediated locking of Tyr^7.53^ in one single conformation, which was fully released only upon Asp^2.50^-protonation (Fig. 4D), prohibited Tyr^7.53^ from coupling its dynamics to other residues and thereby sharing information.

**Fig. 5.**
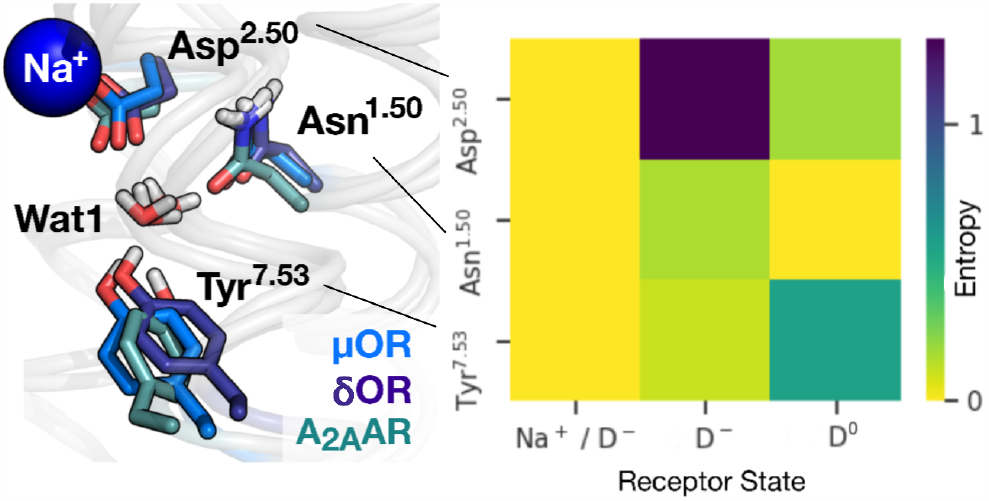
Sodium confines Asp^2.50^, Asn^1.50^ and Tyr^7.53^ to single conformations, inhibiting communication among residue networks. Exemplar frames from molecular dynamics simulations of *µ*OR, δOR and A_2A_AR in the Na^+^/D^-^ simulation state (left). The side chain conformational state entropy (right) of the sodiumbound conformations of Asp^2.50^, Asn^1.50^ and Tyr^7.53^ is decreased to zero in all three receptors, fixing each residue to a specific conformation.

Additionally, Pro^5.50^ of the P^5.50^-I-F motif exhibited zero entropy, i.e. confinement to a single conformation, for the sodium-bound state. As opposed to protonation, SSI further indicated that sodium expulsion exerted an influence on both I^3.40^ and F^6.44^ of this motif. This highlights that sodium interacts with the entire P-I-F motif, which is known to be strongly coupled to ligand binding (23), as F^6.44^ directly interacts with W^6.48^, forming the base of the extracellular ligand binding pocket (47). By contrast, all microswitches displayed fluctuations between various different conformational states in the sodium-free (D^-^) simulations. In the protonated Asp^2.50^ state, only Asn^1.50^ was restrained to a single conformation in its vicinity.

In summary, we conclude that the sodium ion holds the receptor in the inactive state by restraining the key microswitches Asp^2.50^, Tyr^7.53^, Asn^1.50^, and Pro^5.50^ to single conformations. As they are locked in specific states, these microswitches are unable to participate in information transfer, disconnecting the information transfer route across the receptors that emerges fully only upon Asp^2.50^ protonation. Since sodium dissociation is strongly coupled to Asp^2.50^ protonation (12), the sodium–Asp^2.50^ pair forms a single interlinked switch, governing information flow in GPCRs.

## Discussion

Ions play important roles in GPCR signal transduction, affecting both their signal bias (48) and receptor activation (3, 4, 6, 12). In particular, a sodium ion bound to a highly conserved ionizable binding site (3, 6, 7, 11, 24) is known to play a key part in activating class A GPCRs. Signal transmission is further thought to be underpinned by a tightly connected network of polar residues and internal water molecules that extends across the receptors (14, 17) and by the conformational changes of protein microswitches (1, 16, 23). How these components interact to affect activation and signaling, however, has been unclear. Here, we applied information theory to *µ*s-timescale simulations of three class A GPCRs to identify coupled transitions between these elements and examine their relation to receptor activation. Our results revealed that a long-range information transfer pathway connects the most highly-conserved protonatable residues in class A GPCRs (Asp^2.50^, Arg^3.50^ and Asp^3.49^) from the extracellular ion binding site to the intracellular G-protein interface (Fig. 6). The connectivity was established by protonation of Asp^2.50^, triggered by the removal of a sodium ion that is known to bind to Asp^2.50^ in the inactive receptor state (1, 4).

**Fig. 6.**
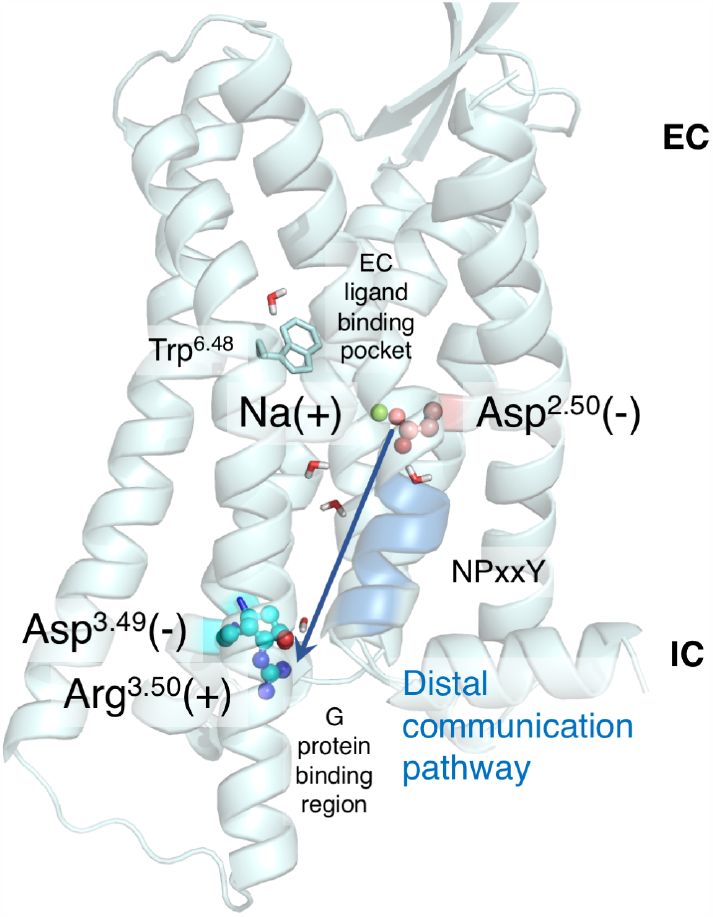
A long-range information transfer pathway connects the conserved ionizable residues Asp^2.50^, Arg^3.50^ and Asp^3.49^, from the extracellular ion binding site to the intracellular G-protein interface. This allows protonatable sites at the G-protein interface regions to receive information about the Asp^2.50^ protonation state, and implicates Asp^2.50^ protonation in G-protein binding. EC, extracellular; IC, intracellular.

A network of ionizable residues extending through GPCRs is thought to be involved in GPCR signal transduction and may represent an evolutionary link to microbial rhodopsins (9, 12, 24, 49, 50). Our findings therefore suggest that the long-range information pathway we identified may couple the protonation states of the distal conserved Asp^2.50^ and Arg^3.50^/Asp^3.49^.

In the sodium-independent, visual class A GPCR rhodopsin, two protonation switches are known to initiate activation. The first protonation event is associated with the light-induced transition of the retinal Schiff base, central to the receptor similar to Asp^2.50^ in non-visual receptors, while the additional protonation occurs at the D^3.49^RY motif (51, 52). The coordinated action of such a conserved twin-protonation switch during activation of class A GPCRs would likely require a mechanism to relay protons, and exchange information about the states of the two distal sites. The information pathway we found between Asp^2.50^ and Asp^3.49^ is composed of polarizable water molecules and residues with conformations that couple to hydrogen bonding patterns at Asp^2.50^, and therefore well suited to underpin proton transfer.

In many cases, ligand binding alone is thought to provide insufficient energy for receptors to transition between inactive and active conformations (3, 9). However, GPCRs are able to harness energy from the membrane potential gradient, potentially through ion and proton transfer (9, 12, 50, 53). Even in the presence of bound antagonists, we observed conformational rearrangements within highly conserved microswitches in the DRY and NPxxY motifs upon protonation of three receptors, which were reminiscent of the transition between inactive and active state crystal structures. This stochastic, ligand-independent sampling of active state conformations is in line with reports that suggest spontaneous Asp^2.50^-protonation and egress of the sodium ion may be linked to basal signalling (12), as well as the observation that active state structures exhibit no bound sodium (4). Our results suggest that sodium unbinding and protonation events, potentially coordinated between multiple conserved protonatable sites, play a key role in the activation process of class A receptors.

Two water molecules, structurally conserved across both active and inactive receptor states, are known to facilitate water-mediated interactions that correlate with the inactive (water 1) and active (water 2) receptor states (14). We found that these two water molecules function as microswitches which couple to the ion binding states of Asp^2.50^. Both waters are essential for the transmission of information along a conserved pathway between the ion binding site and the DRY motif. Other water molecules play further important roles in connecting protein microswitch states over long-ranges. By disabling protonation and tightly restraining the conformation of Asp^2.50^ and neighboring residues and waters, sodium stabilised the inactive state of the receptor and disconnected these information transfer pathways.

Only a small fraction of class A GPCRs do not possess a sodium binding site in their transmembrane domain (for example the NK1 neurokinin receptor (4, 54)), however they contain a similarly dense internal network of polar residues and water. Whereas Asp^2.50^ is replaced by Glu, the DRY motif is still present in these receptors. A sodium-independent mechanism of switching protonation states and rearranging the polar network is conceivable in these cases. However, future studies will be necessary to confirm if the polar signal transmission and activation mechanism we find is conserved in the limited number of atypical class A GPCRs.

## Materials and Methods

### Defining Microswitch States

Residue side chain rotamer conformations were defined by in-house Gaussian style clustering to *χ*1 and *χ*2 dihedral angles, as seen in Fig. 7. Rotamer angles were determined every 240 ps using gmx_chi. The frame separation was chosen to be slightly smaller than the autocorrelation time of the conserved water dynamics with the fastest relaxation time (55). A Hanning window function was used to smooth the probability density function to remove noise and locate the state maxima for each rotamer conformation. Guess parameters for Gaussian fitting were then obtained by locating the full-width at half-maximum from each maximum and Gaussian curve fitting was applied. The microswitch conformational state limits were defined as the Gaussian intersects. Each discretized distribution was additionally checked visually to ensure the fitting was accurate.

**Fig. 7.**
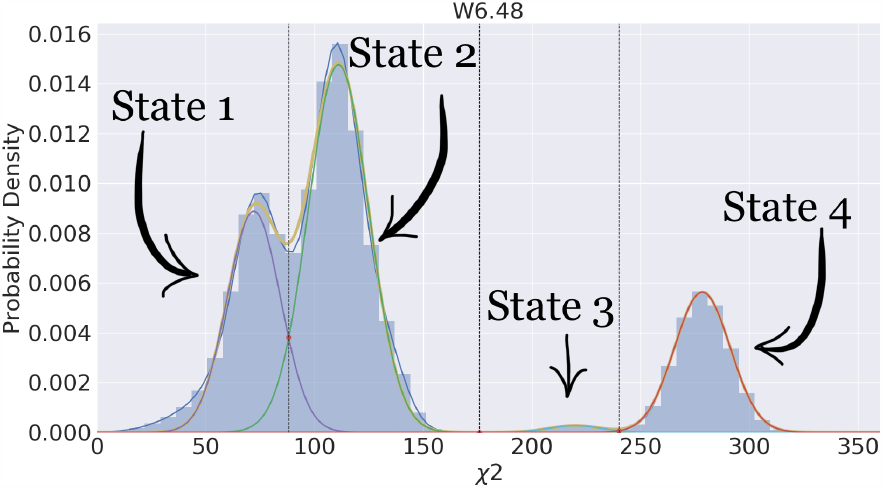
Illustration of the Gaussian clustering method employed for defining discrete residue rotamer states. The example given is for the *χ*2 angle of the highly conserved tryptophan residue Trp^6.48^. State limits are defined as the intersects of each Gaussian and the maximum and minimum values of the angular distribution.

Water pocket states were defined by water molecule occupancy and using the water dipole moment to determine the molecular orientation. For *µ*OR, water pockets were defined using MDAnalysis (56) as spheres of radius 4 Å centred on the geometric centres of (i) the tetrahedron formed by joining the C-*α* atoms of Asn^1.50^-Asp^2.50^-Asn^7.49^-Leu^2.46^; (ii) the triangle formed by joining the C-*α* atoms of Asn^7.45^-Asn^7.49^-Ala^6.38^; (iii) the triangle formed by joining the C-*α* atoms of Cys^6.47^-Pro^6.50^-Cys^7.37^; and (iv,v) spheres of 5 Å radius centred on the geometric centres of the tetrahedron formed by joining the C-*α* atoms of Asn^2.45^-Thr^3.42^-Val^4.45^-Trp^4.50^, and the triangle Val^3.48^-Arg^34.57^-Ala^4.42^, respectively, as seen in Fig. 8. Similar positions were used for the *δ*OR and A_2A_AR, determined by centering the water probability density in the geometric centre of the selected C-*α* atoms.

**Fig. 8.**
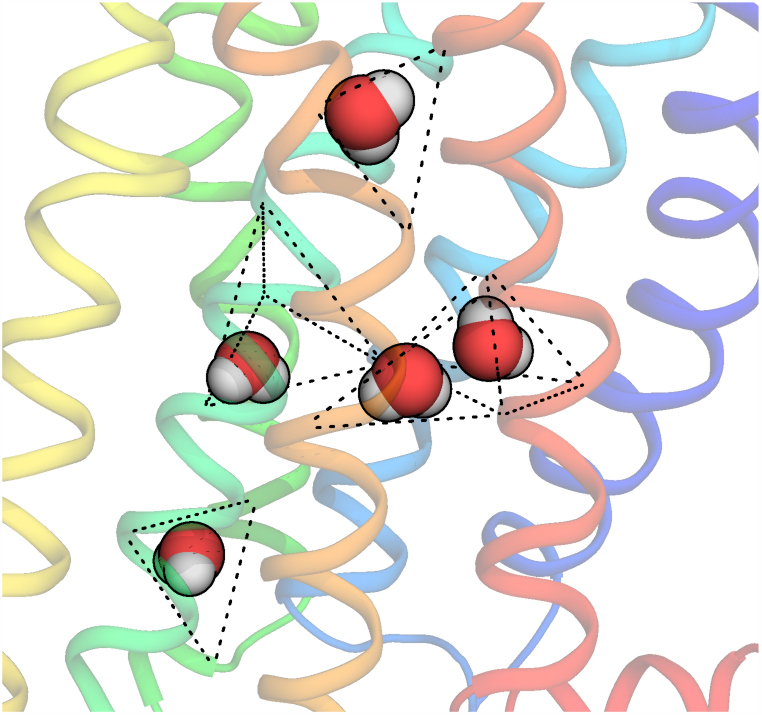
Water pockets conserved in active and inactive crystal structures of *µ*OR, A_2A_AR and *δ*OR. Pockets are defined as spheres of radii 4 Å and 5 Å centred on the centre of geometry of the respective triangles and tetrahedrons.

When water was not found in a specific water pocket at a certain time point, that pocket was assigned a discrete empty state. Water pockets must have all three water atoms within the pocket as defined above to be considered occupied. When water was found inside a pocket, the dipole moment vector for the water molecule 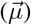 was calculated by 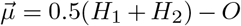, where *H*_*i*_ is the (x,y,z) coordinate of the *i*th hydrogen, and *O* is the (x,y,z) coordinate of the oxygen. The orientation of water molecules was then calculated by converting the dot product of the dipole vector and the simulation box axes vector into spherical coordinates, 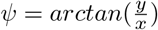 and 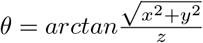, followed by the same Gaussian style clustering algorithm. Together, this allowed us to deduce the functional effect of each water pocket by combining the continuous distributions of water polarisation and the binary/discrete distributions of water pocket occupancy (i.e. occupied vs. unoccupied) into a univariate distribution.

#### Information Theory

Unlike cartesian correlation approaches, a state based mutual information approach resolves slow modes in protein dynamics that have the greatest functional relevance, reducing noise levels (27, 39). Causality can be inferred from our state based calculation of mutual information because the ion binding states in each ensemble are constrained (i.e. they are constant for each simulation), only changing state upon concatenation, and therefore the root change occurs at the ion binding site. The information transfer resolved therefore travels from the ion binding site to residues and internal waters. However, since SSI is symmetric in principle, note that the identified information pathway could also serve to support information flow towards the ion binding site.

The conformational state entropy (Shannon entropy) of microswitch *X* was determined from *H*(*X*) = − ∑_*i*_ *p*(*i*)*log*_2_*p*(*i*), where *p*(*i*) is the probability of state *i* occurring, defined using our definition of discrete states (Fig. 7). *H*(*X*) is also a measure of the maximum information a microswitch can store (26). A regular average was taken across all common Ballesteros-Weinstein positions in the three receptors to deduce the average entropy of a specific residue. We used the state limits of the concatenated simulations for the conformational state entropy to extrapolate conclusions about the SSI transfer, thereby identifying the whether a microswitch is prohibited from communicating in a specific simulation state. Therefore, residues have zero entropy only if they occupy one of the information transfer states, and not just a single Gaussian distribution that spanned two information states.

SSI (mutual state information) between two microswitches *X* and *Y* was calculated using *SSI*(*X, Y*) = *H*(*X*) + *H*(*Y*) − *H*(*X, Y*), where *H*(*X, Y*) is the joint entropy, quantifying the entropy of individual microswitches *X* and *Y* as one multivariate microswitch using our definition of discrete states (26).

The co-SSI (or interaction information) was calculated using

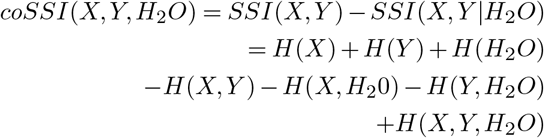

where *SSI*(*X, Y*| *H*_2_*O*) determines what magnitude of SSI shared between *X* and *Y* is dependent on the state of a specific third component, often an intercalating water molecule (*H*_2_*O*) (29). There are different interpretations of negative interaction information values in the literature (30–32). We adopted the interpretation used by LeVine & Weinstein, in which negative interaction information serves to attenuate information transfer between two residues (31). We deemed SSI and co-SSI values of less than 0.05 bits insignificant on a 95% significance level. As the changes made to the state of the ion binding site were binary (i.e. 1 bit of information – sodium-bound to sodium-free state; and sodium-free to /Asp^2.50^-protonated state), the theoretical upper limit for all SSI and co-SSI values was therefore 1 bit.

To focus on information transfer pathways conserved in all three receptors, we determined the geometric mean of SSI and co-SSI from all common residue Ballesteros-Weinstein positions.

#### Protein structure preparations

All crystal structures (*µ*OR - pdb:4dkl, *δ*OR - pdb:4n6h, A_2A_AR - pdb:5olz) were obtained from GPCRdb (57), selecting the GPCRdb refined structure in which mutations are reversed to the canonical sequence and cleaved loops are re-modelled using MODELLER 9.18 to produce the final structure (58, 59). All transmembrane crystalline waters and ligands were kept in the structure, removing all external waters and non-protein molecules. The proteins were truncated and capped with acetyl and methyl groups at corresponding N and C terminals using PyMOL (60). As the *µ*OR crystal structure did not resolve a sodium ion in the pocket, the sodium-bound structure was obtained by aligning the *δ*OR crystal structure and placing the sodium ion in an identical position. Asp^2.50^-charged protein structures were prepared by removing sodium from the crystal structure and ensuring no sodium ion re-associated during the course of the simulation. The third, protonated-Asp^2.50^ protein state was prepared in the molecular dynamics software GROMACS-5.1.1 (61). Ligand structures were taken from their respective protein pdb files, and parameterised for molecular dynamics simulations in GROMACS-5.1.1 using ACPYPE (62).

#### MD simulations

We modelled the protein, membrane, and ligand using the amber99sb-ildn forcefield (63) in GROMACS-5.1.1 with virtual sites to allow a time-step of 4 fs. The proteins were embedded in a pre-equilibrated SLipid POPC membrane (64) using InflateGRO (65). The protein-membrane complexes were solvated with a neutral solution of TIP3P water molecules (66) containing NaCl at ∼150 mM concentration, with a box size of ∼128×134×125 Å^3^. The systems were equilibrated in both the NVT and NPT ensembles at 310 K for 3 ns with position restraints on the ligand-receptor complex heavy atoms, with a further 70 ns of production run simulation considered as additional NPT equilibration. Following equilibration, simulations were performed for 1.7 *µ*s each at a constant temperature of 310 K and pressure of 1 bar, with the protein-ligand complex, membrane, and solution independently coupled to a temperature bath using a Nosé-Hoover thermostat with a time constant of 0.5 ps and a semi-isotropic Parrinello-Rahman barostat with a time constant of 5 ps (67). The trajectory of the *δ*OR protonated at Asp^2.50^ was obtained from a previous simulation study (53, 68). All protein and lipid bond lengths were constrained with the LINCS algorithm (69), while water bond lengths were constrained using SETTLE (70).

## SUPPLEMENTARY MATERIALS

Fig. S1: Asp^2.50^-protonation SSI.

Fig. S2: Sodium Expulsion SSI.

Fig. S3: Inter-hydroxyl distance between Tyr5.58-Tyr7.53 of *µ*OR.

Fig. S4: Activation coordinate (TM2-TM6) in the *δ*OR.

Fig. S5: Asp^2.50^-protonation co-SSI For Wat1.

Fig. S6: Asp^2.50^-protonation co-SSI For Wat2.

Fig. S7: Asp^2.50^-protonation co-SSI For Wat3.

Fig. S8: Asp^2.50^-protonation co-SSI For Wat4.

Fig. S9: Asp^2.50^-protonation co-SSI For Wat5.

Fig. S10: Sodium Expulsion co-SSI For Wat1.

Fig. S11: Sodium Expulsion co-SSI For Wat2.

Fig. S12: Sodium Expulsion co-SSI For Wat3.

Fig. S13: Sodium Expulsion co-SSI For Wat4.

Fig. S14: Sodium Expulsion co-SSI For Wat5.

## ACKNOWLEDGEMENTS

We thank Seva Katritch, Andrei Pisliakov, and Martin Vögele for critical reading of and comments on the manuscript.

## AUTHOR CONTRIBUTIONS

Conceptualization: N.J.T and U.Z. Methodology: N.J.T and U.Z. Formal analysis: N.J.T and U.Z. Investigation: N.J.T, O.N.V and U.Z. Writing (original draft): N.J.T and U.Z. Writing (review and editing): N.J.T, C.M.I and U.Z. Supervision: U.Z.

