## Supplementary Information for "Ion-water coupling controls class A GPCR signal transduction pathways"

**Asp$^{2.50}$-protonation SSI**

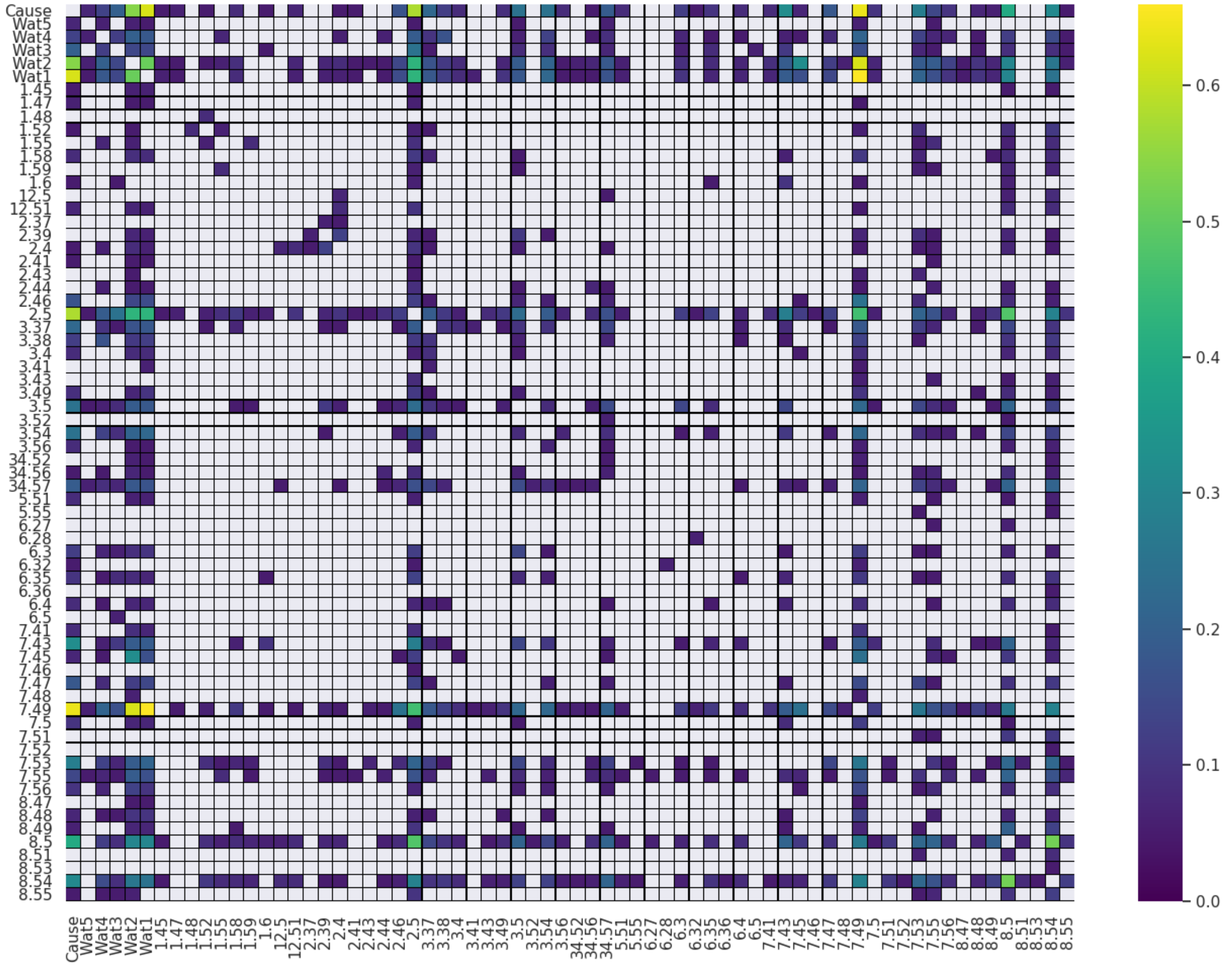


Fig. S 1: **Matrix of SSI values for Asp$^{2.50}$ protonation of the A$_{2A}$-adenosine, $\delta$-opioid, and $\mu$-opioid receptors (bits; geometric means).** We refer to the information source, i.e. Asp$^{2.50}$-protonation, as 'cause'. SSI values of zero are masked from the SSI matrix, therefore appear as white cells in the heatplot. Here we only include microswitches that share significant SSI with at least one other microswitch in the heatplot, with all additional microswitches sharing zero total SSI.

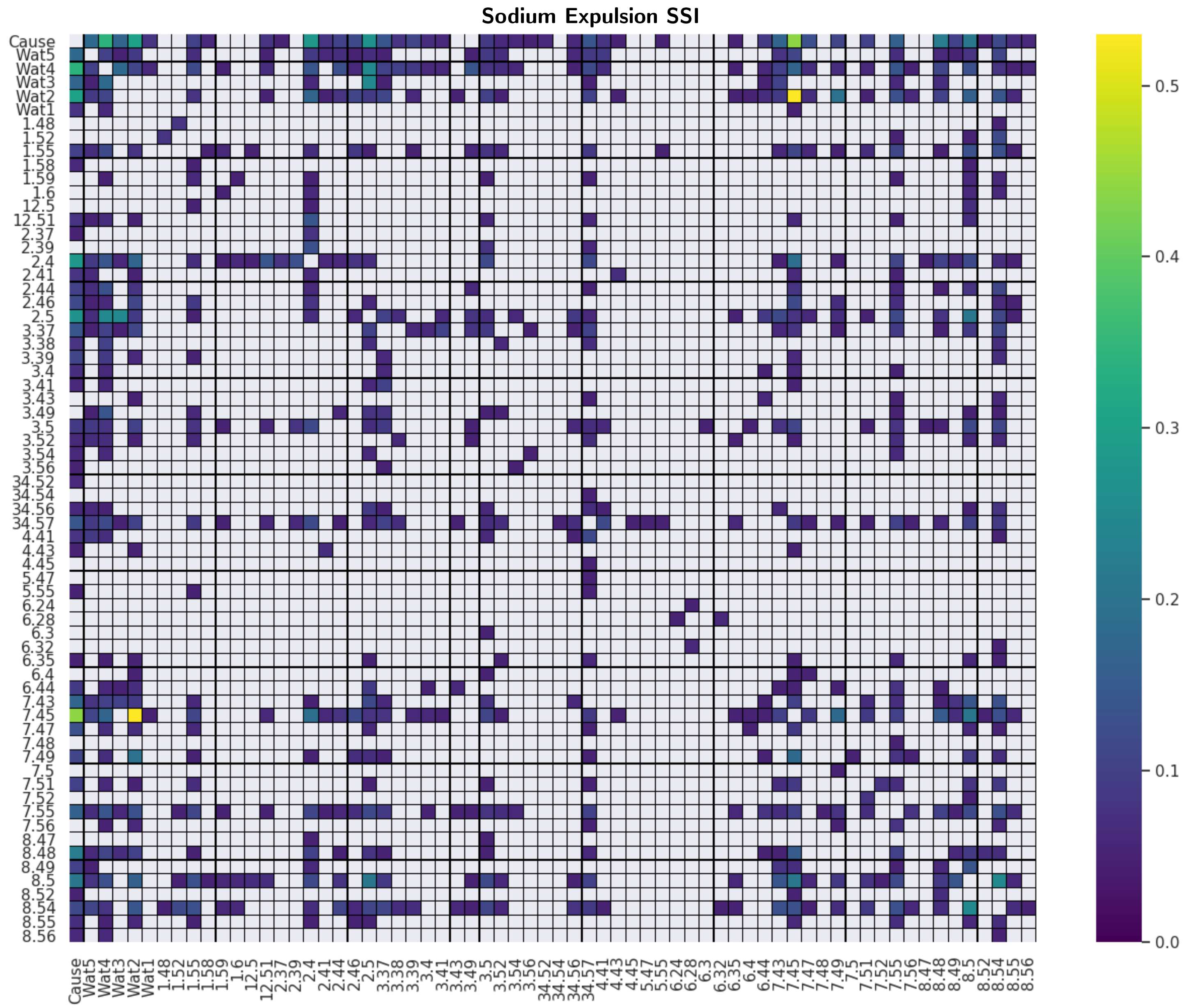


Fig. S 2: **Matrix of SSI values for sodium removal in the $A_{2A}$-adenosine, $\delta$-opioid, and $\mu$-opioid receptors (bits; geometric means).** We refer to the information source, i.e. sodium expulsion, as 'cause'. SSI values of zero are masked from the SSI matrix, therefore appear as white cells in the heatplot. Here we only include microswitches that share significant SSI with at least one other microswitch in the heatplot, with all additional microswitches sharing zero total SSI.

### Inter-hydroxyl distance between $Tyr^{5.58}$-$Tyr^{7.53}$ of $\mu$OR.

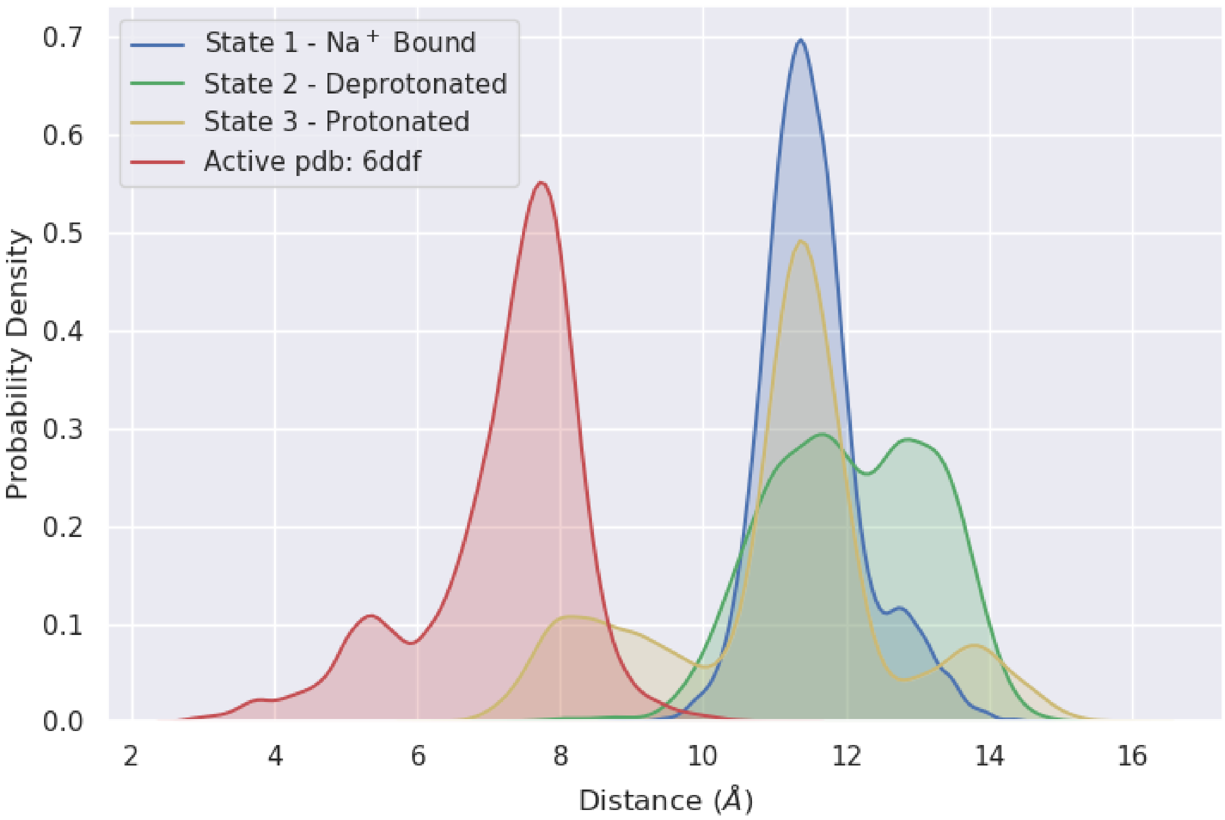


Fig. S 3: **Inter-hydroxyl distance between $Tyr^{5.58}$-$Tyr^{7.53}$ of $\mu$OR.** Probability distribution for the distance between OH-groups of $Tyr^{7.53}$ and $Tyr^{5.58}$ throughout the three receptor states $Na^{+}/D^{-}$ (state 1), $D^{-}$ (state 2), and $D^{0}$ (state 3) for $\mu$OR (pdb: 4dkl), and the active state $\mu$OR structure (pdb: 6ddf). The $D^{0}$ simulation samples the decreased distance seen in the active-state distribution, representing the downward flip of $Tyr^{7.53}$ correlated with opening of the hydrophobic barrier.

**Activation coordinate (TM2-TM6) in the $\delta$OR.**

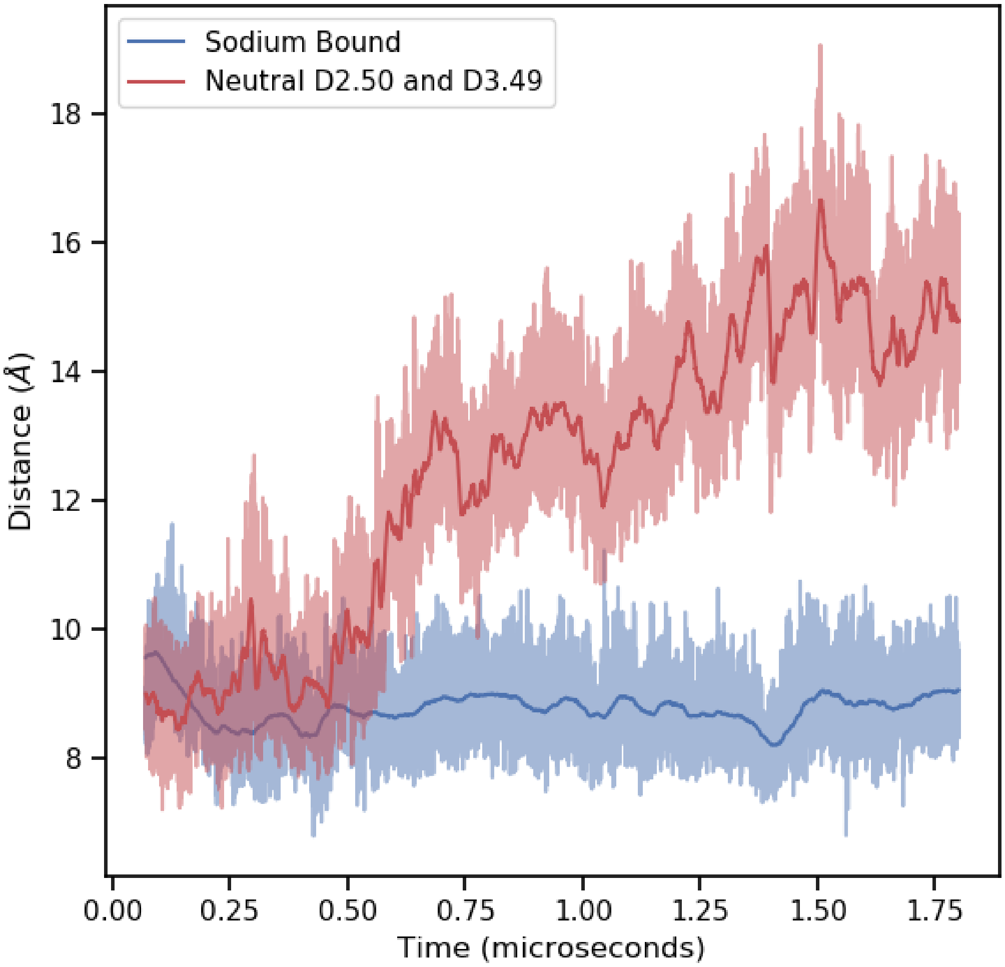


Fig. S 4: **Activation coordinate (TM2-TM6) in the $\delta$OR.** The distance between Thr$^{2.39}$-C$\alpha$ and Ile$^{6.33}$-C$\alpha$ for the sodium bound, charged Asp$^{2.50}$ and Asp$^{3.49}$ receptor simulation (blue line) and a sodium not bound, protonated-Asp$^{2.50}$ and protonated-Asp$^{3.49}$ simulation (red line). The increase in distance seen for the doubly-protonated simulation represents the opening of the intra-cellular effector binding site seen upon activation.

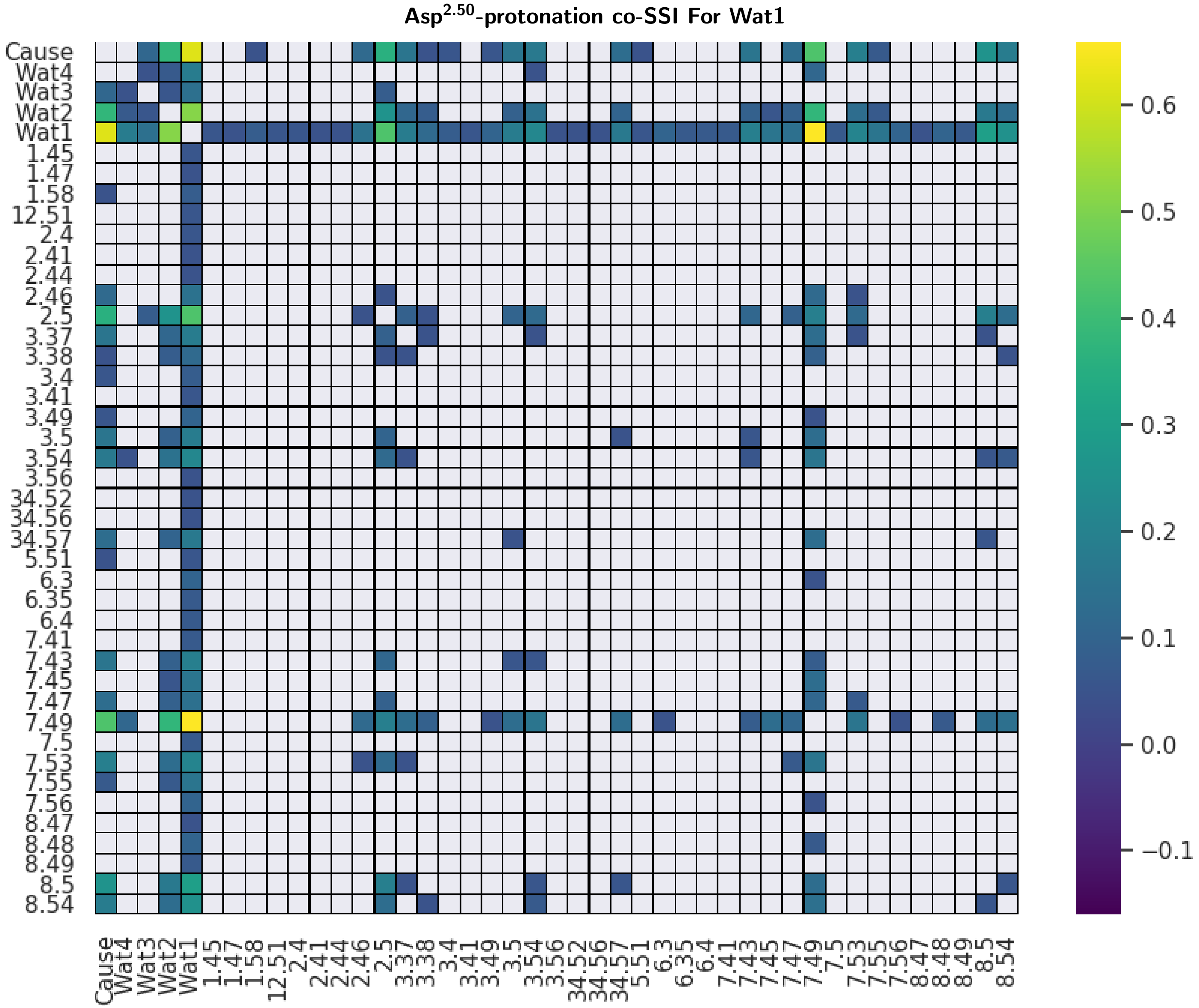


Fig. S 5: **Matrix of co-SSI values showing impact of water 1 (wat1) on SSI transfer between all microswitch pairs for Asp$^{2.50}$-protonation in the A$_{2A}$-adenosine, $\delta$-opioid, and $\mu$-opioid receptors (bits; geometric means).** We refer to the information source, i.e. Asp$^{2.50}$-protonation, as 'cause'. Co-SSI values of zero are masked from the co-SSI matrix, therefore appear as white cells in the heatplot. Here we only include microswitches that share significant co-SSI with at least one other microswitch in the heatplot, with all additional microswitches sharing zero total co-SSI.

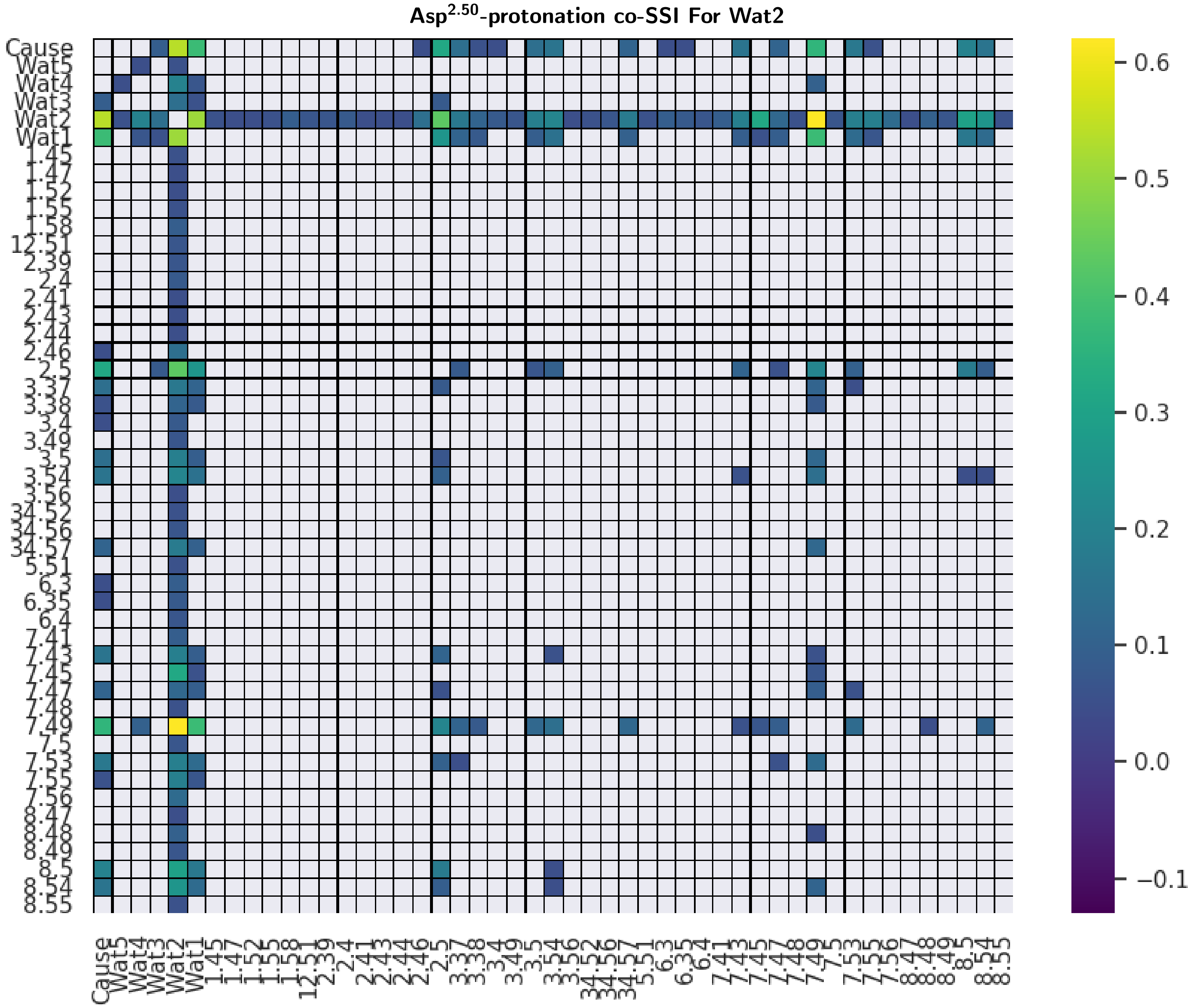


Fig. S 6: **Matrix of co-SSI values showing impact of water 2 (wat2) on SSI transfer between all microswitch pairs for Asp$^{2.50}$-protonation in the A$_{2A}$-adenosine, $\delta$-opioid, and $\mu$-opioid receptors (bits; geometric means).** We refer to the information source, i.e. Asp$^{2.50}$-protonation, as 'cause'. Co-SSI values of zero are masked from the co-SSI matrix, therefore appear as white cells in the heatplot. Here we only include microswitches that share significant co-SSI with at least one other microswitch in the heatplot, with all additional microswitches sharing zero total co-SSI.

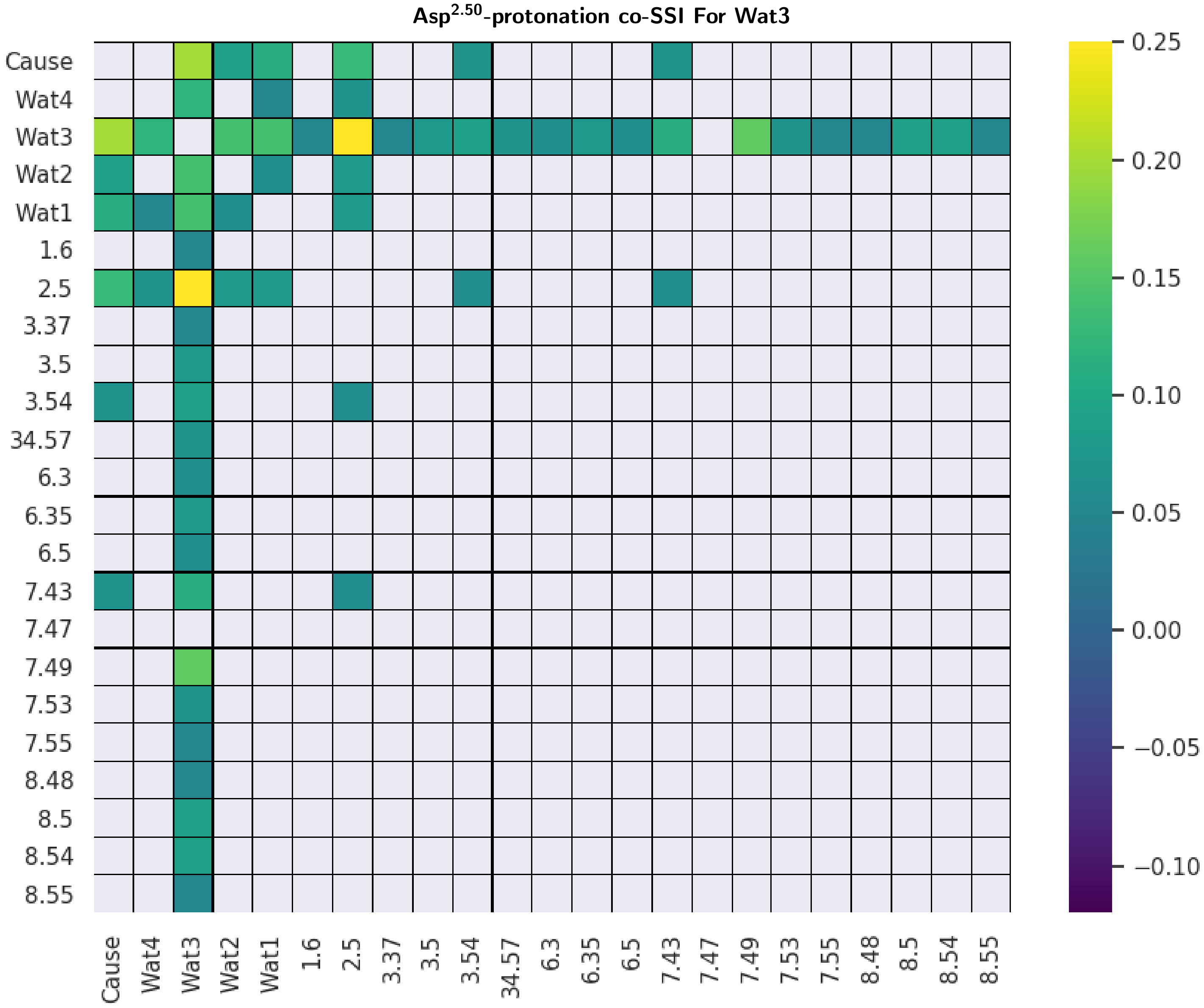


Fig. S 7: **Matrix of co-SSI values showing impact of water 3 (wat3) on SSI transfer between all microswitch pairs for Asp$^{2.50}$-protonation in the $A_{2A}$-adenosine, $\delta$-opioid, and $\mu$-opioid receptors (bits; geometric means).** We refer to the information source, i.e. Asp$^{2.50}$-protonation, as 'cause'. Co-SSI values of zero are masked from the co-SSI matrix, therefore appear as white cells in the heatplot. Here we only include microswitches that share significant co-SSI with at least one other microswitch in the heatplot, with all additional microswitches sharing zero total co-SSI.

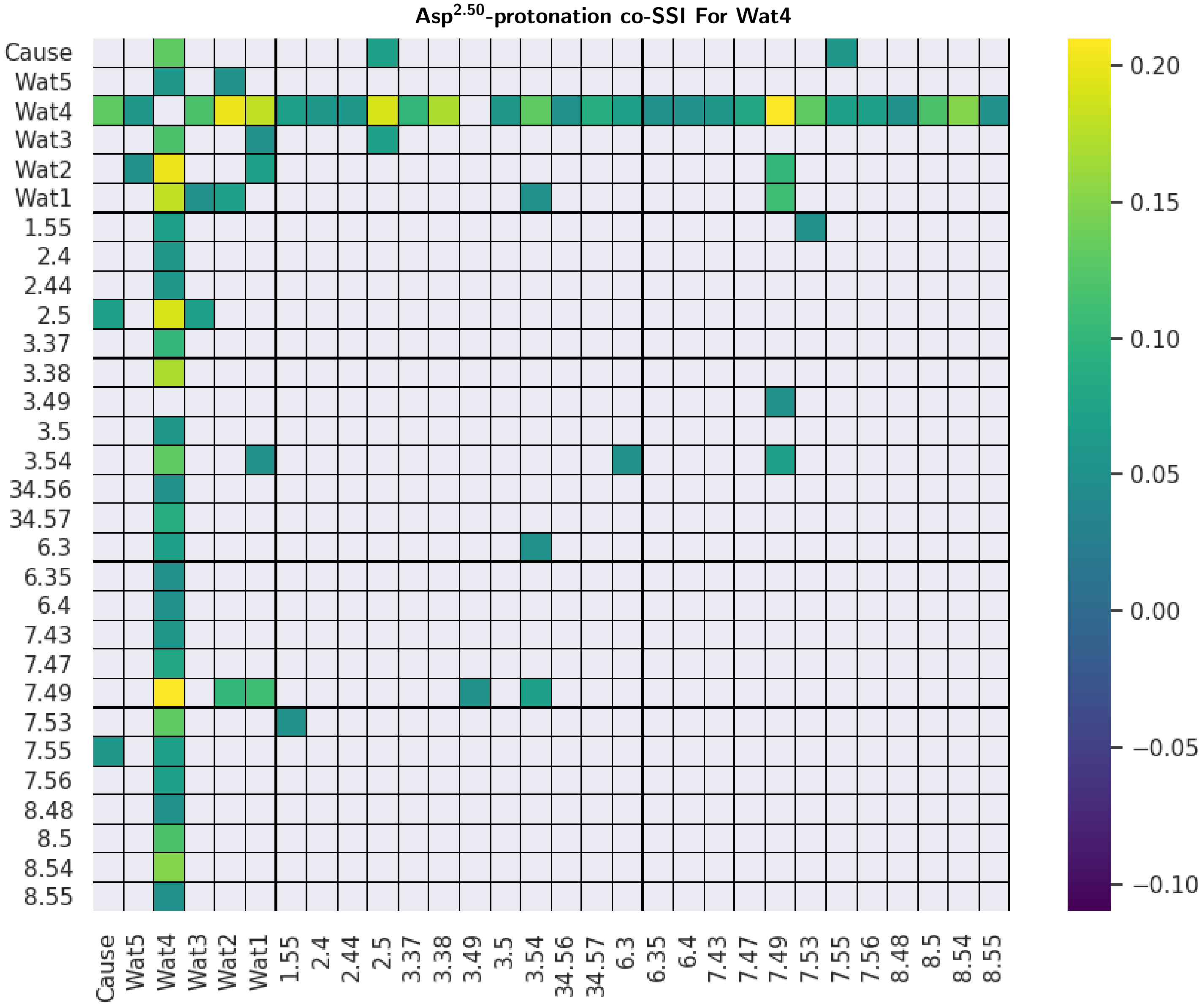


Fig. S 8: **Matrix of co-SSI values showing impact of water 4 (wat4) on SSI transfer between all microswitch pairs for Asp$^{2.50}$-protonation in the $A_{2A}$-adenosine, $\delta$-opioid, and $\mu$-opioid receptors (bits; geometric means).** We refer to the information source, i.e. Asp$^{2.50}$-protonation, as 'cause'. Co-SSI values of zero are masked from the co-SSI matrix, therefore appear as white cells in the heatplot. Here we only include microswitches that share significant co-SSI with at least one other microswitch in the heatplot, with all additional microswitches sharing zero total co-SSI.

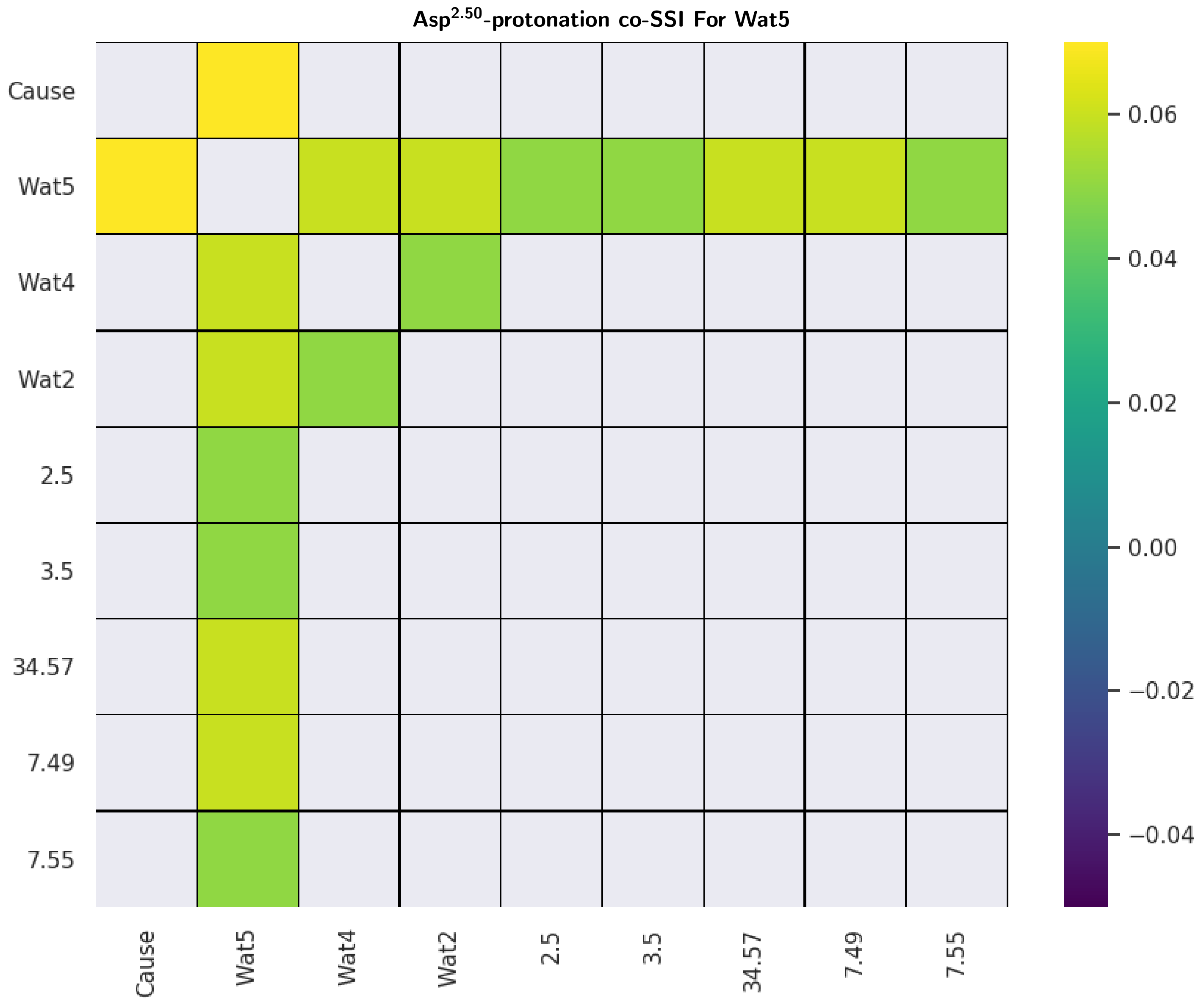


Fig. S 9: **Matrix of co-SSI values showing impact of water 5 (wat5) on SSI transfer between all microswitch pairs for Asp$^{2.50}$-protonation in the $A_{2A}$-adenosine, $\delta$-opioid, and $\mu$-opioid receptors (bits; geometric means).** We refer to the information source, i.e. Asp$^{2.50}$-protonation, as 'cause'. Co-SSI values of zero are masked from the co-SSI matrix, therefore appear as white cells in the heatplot. Here we only include microswitches that share significant co-SSI with at least one other microswitch in the heatplot, with all additional microswitches sharing zero total co-SSI.

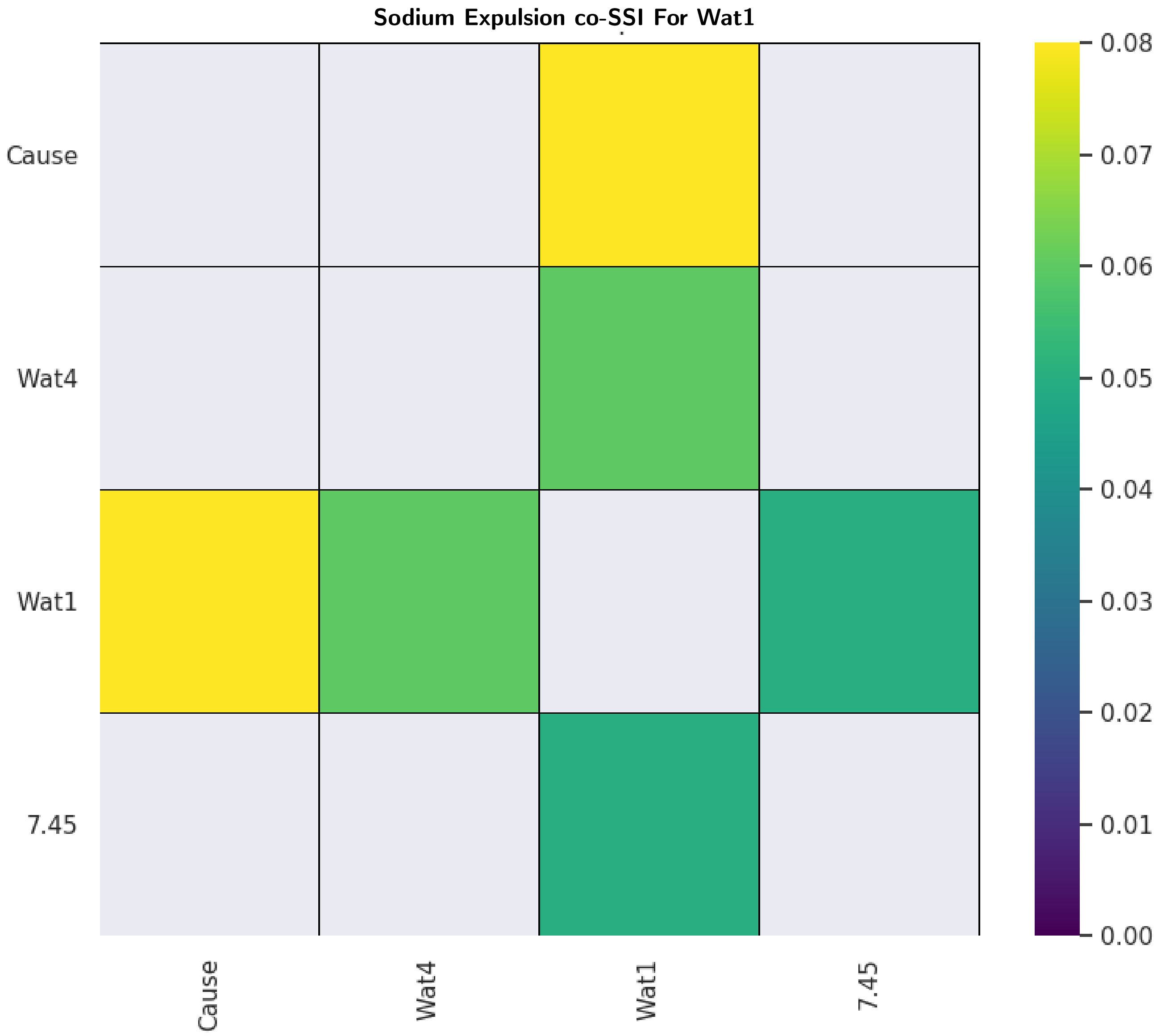


Fig. S 10: **Matrix of co-SSI values showing impact of water 1 (wat1) on SSI transfer between all microswitch pairs for sodium expulsion in the $A_{2A}$-adenosine, $\delta$-opioid, and $\mu$-opioid receptors (bits; geometric means).** We refer to the information source, i.e. sodium expulsion, as 'cause'. Co-SSI values of zero are masked from the co-SSI matrix, therefore appear as white cells in the heatplot. Here we only include microswitches that share significant co-SSI with at least one other microswitch in the heatplot, with all additional microswitches sharing zero total co-SSI.

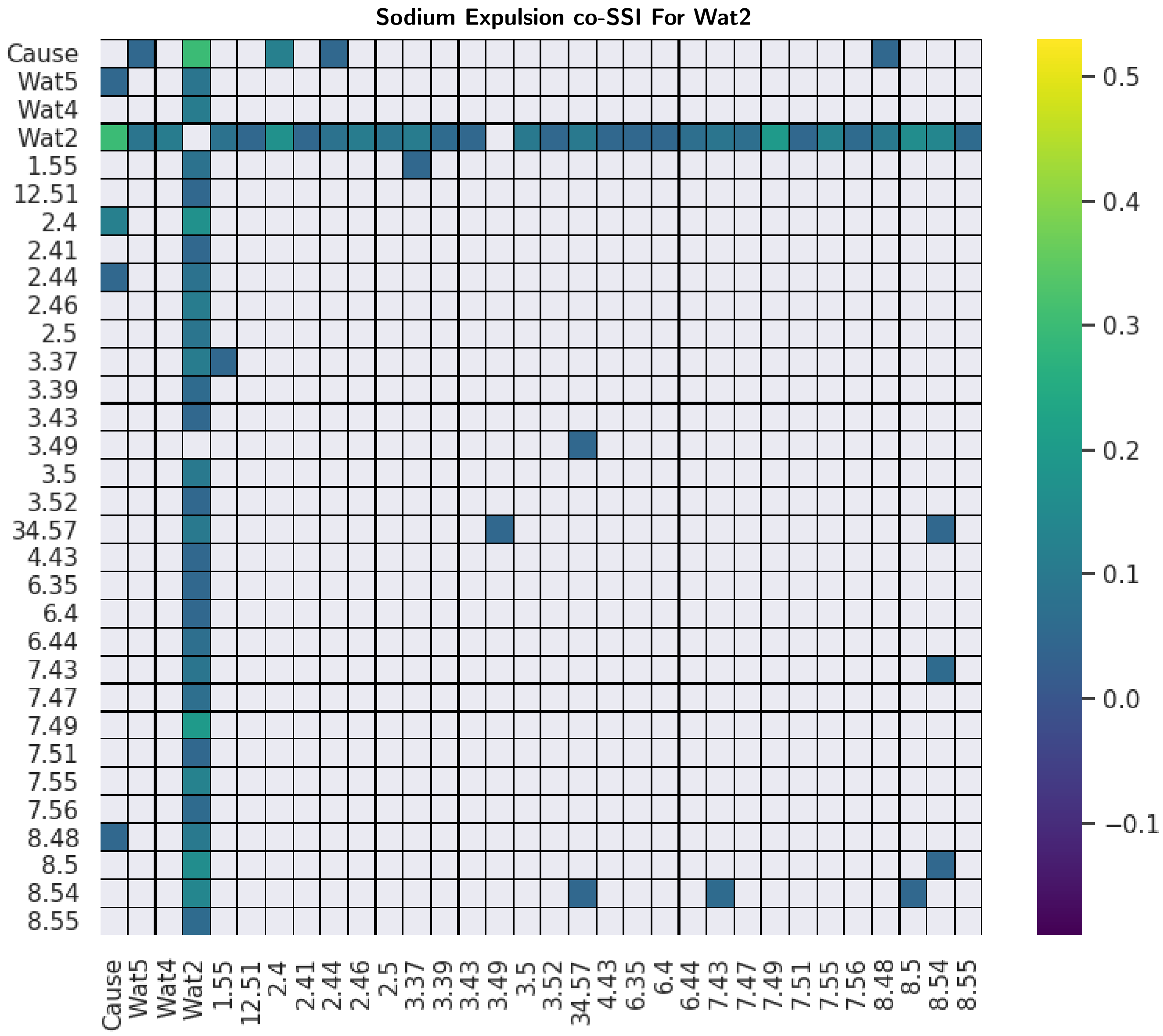


Fig. S 11: **Matrix of co-SSI values showing impact of water 2 (wat2) on SSI transfer between all microswitch pairs for sodium expulsion in the $A_{2A}$-adenosine, $\delta$-opioid, and $\mu$-opioid receptors (bits; geometric means).** We refer to the information source, i.e. sodium expulsion, as 'cause'. Co-SSI values of zero are masked from the co-SSI matrix, therefore appear as white cells in the heatplot. Here we only include microswitches that share significant co-SSI with at least one other microswitch in the heatplot, with all additional microswitches sharing zero total co-SSI.

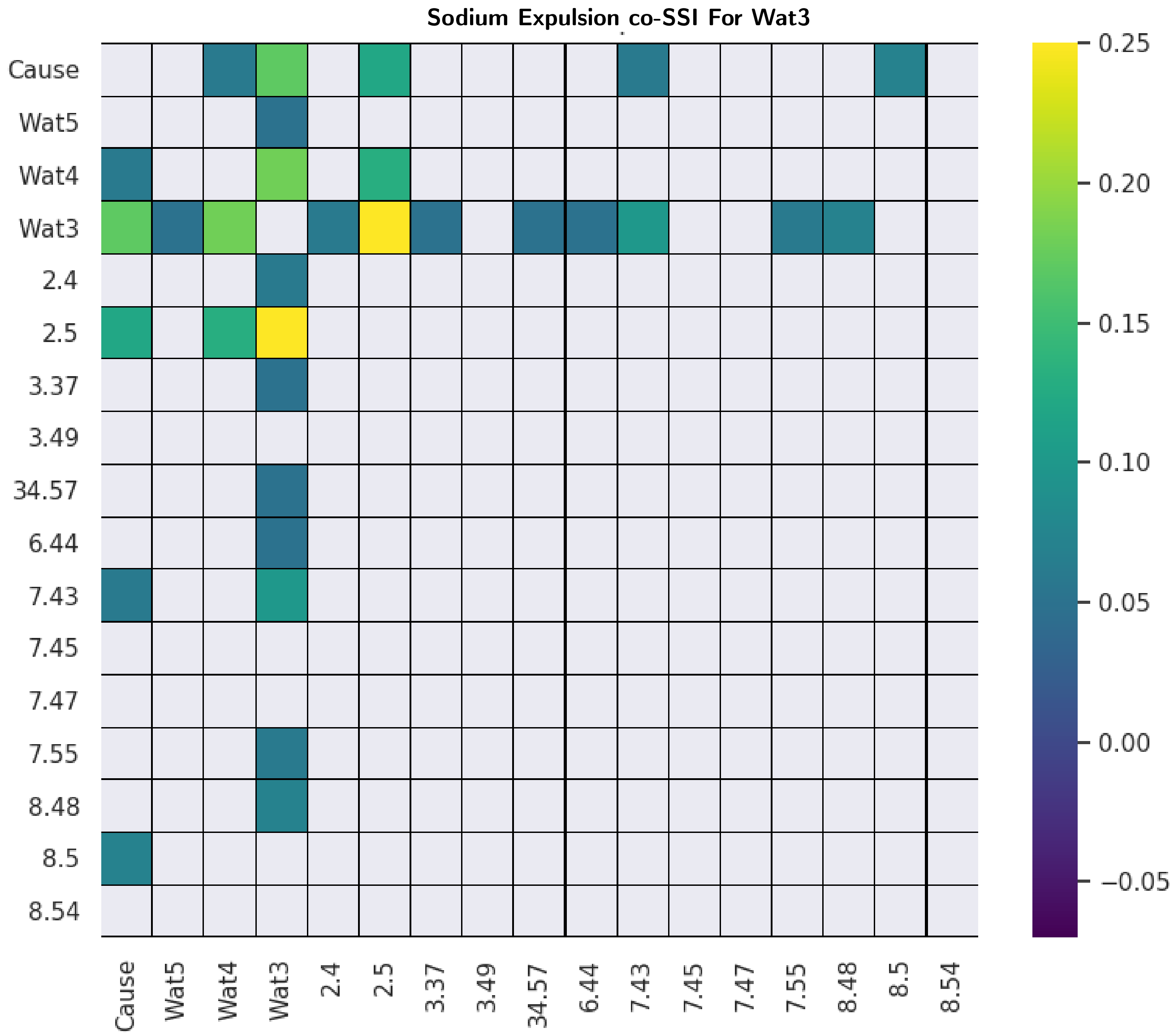


Fig. S 12: **Matrix of co-SSI values showing impact of water 3 (wat3) on SSI transfer between all microswitch pairs for sodium expulsion in the $A_{2A}$-adenosine, $\delta$-opioid, and $\mu$-opioid receptors (bits; geometric means).** We refer to the information source, i.e. sodium expulsion, as 'cause'. Co-SSI values of zero are masked from the co-SSI matrix, therefore appear as white cells in the heatplot. Here we only include microswitches that share significant co-SSI with at least one other microswitch in the heatplot, with all additional microswitches sharing zero total co-SSI.

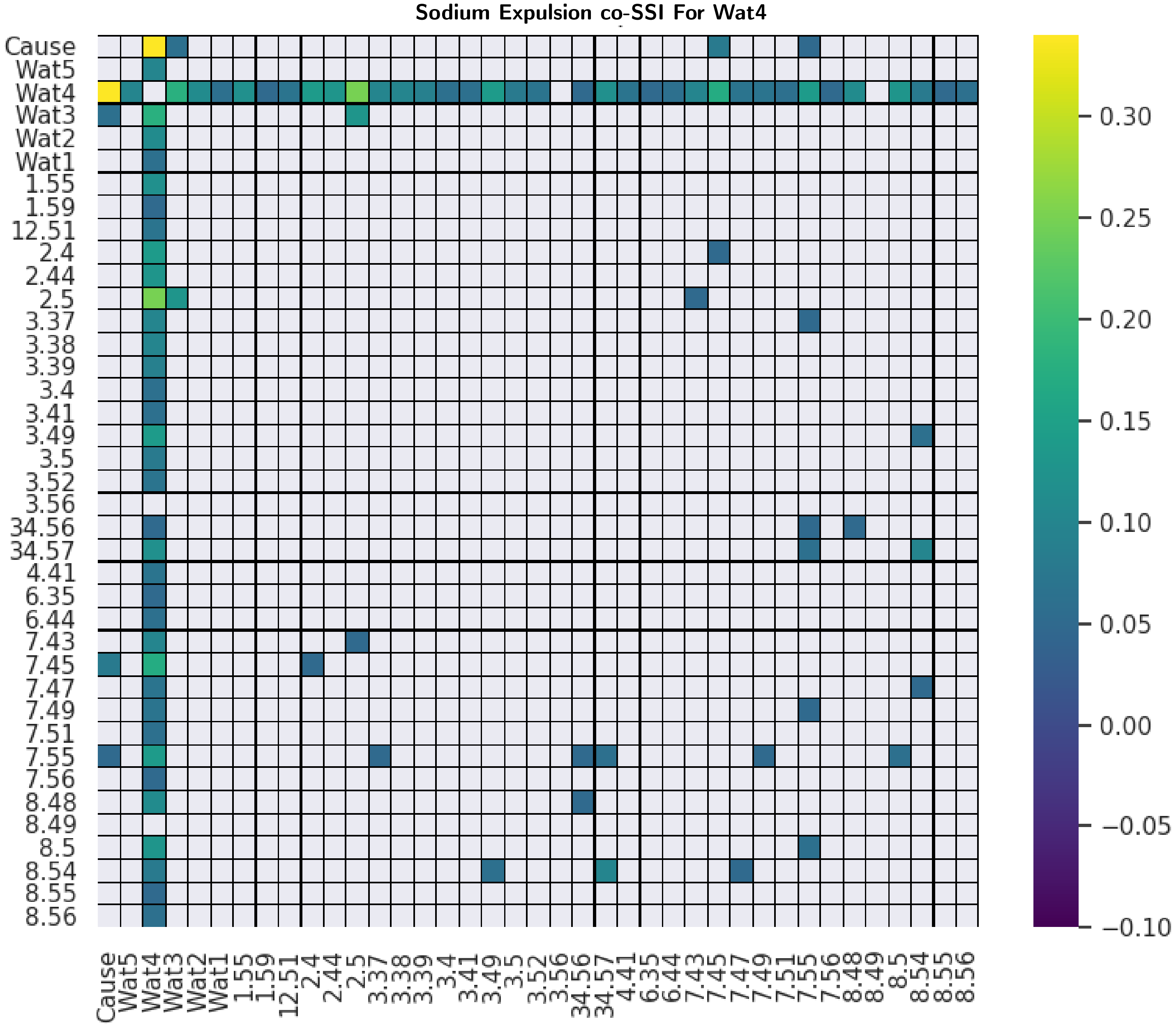


Fig. S 13: **Matrix of co-SSI values showing impact of water 4 (wat4) on SSI transfer between all microswitch pairs for sodium expulsion in the $A_{2A}$-adenosine, $\delta$-opioid, and $\mu$-opioid receptors (bits; geometric means).** We refer to the information source, i.e. sodium expulsion, as 'cause'. Co-SSI values of zero are masked from the co-SSI matrix, therefore appear as white cells in the heatplot. Here we only include microswitches that share significant co-SSI with at least one other microswitch in the heatplot, with all additional microswitches sharing zero total co-SSI.

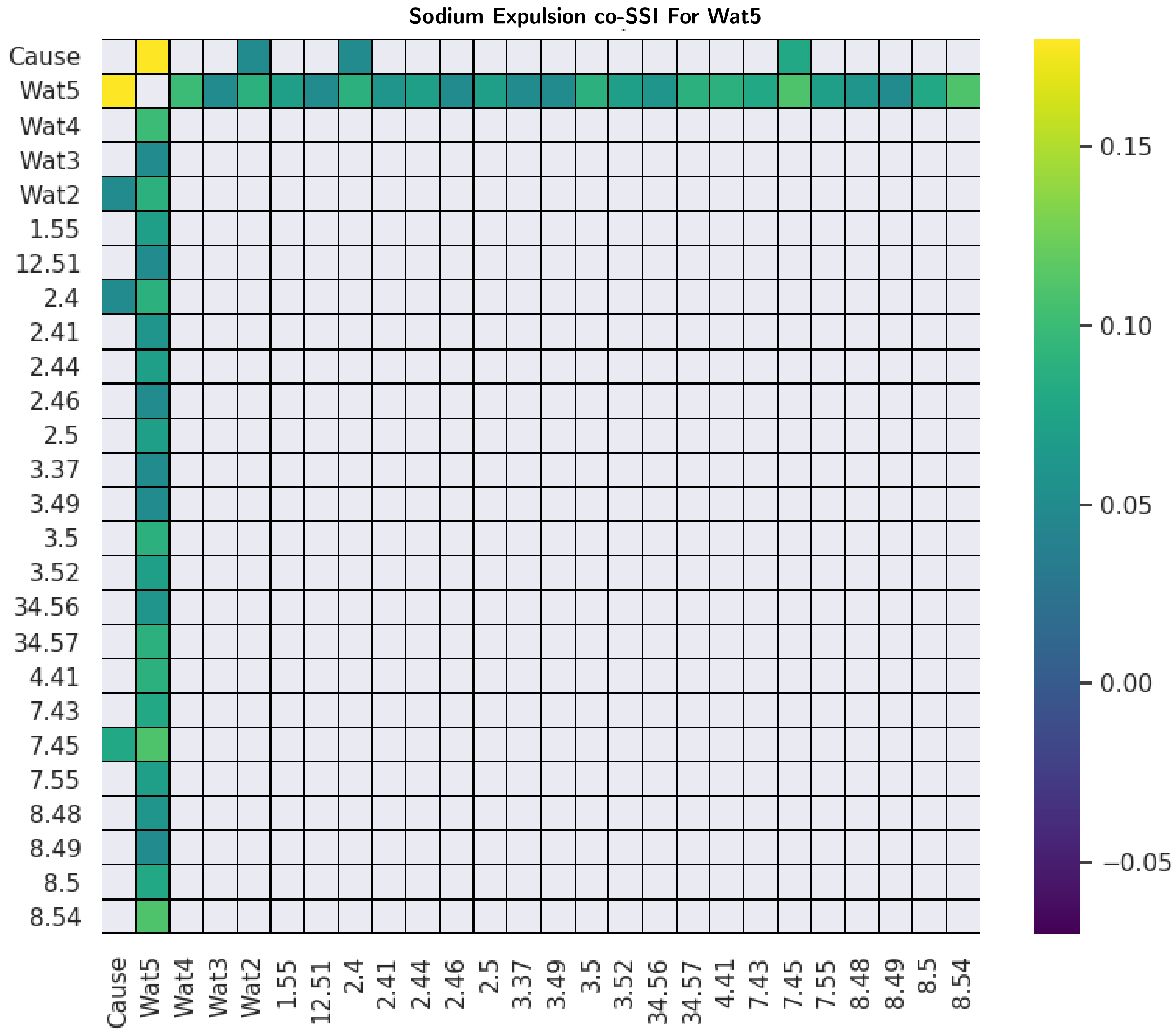


Fig. S 14: **Matrix of co-SSI values showing impact of water 5 (wat5) on SSI transfer between all microswitch pairs for sodium expulsion in the $A_{2A}$-adenosine, $\delta$-opioid, and $\mu$-opioid receptors (bits; geometric means).** We refer to the information source, i.e. sodium expulsion, as 'cause'. Co-SSI values of zero are masked from the co-SSI matrix, therefore appear as white cells in the heatplot. Here we only include microswitches that share significant co-SSI with at least one other microswitch in the heatplot, with all additional microswitches sharing zero total co-SSI.
